# Mitochondrial Hsp60/10 Client Protein Decline Reveals Braak/Tau- and Cognition-Linked Proteostasis Vulnerabilities in Alzheimer’s Disease

**DOI:** 10.64898/2026.06.25.734545

**Authors:** Ashlyn Sloane, Varunya M. Kattunga, Julie K. Andersen

**Author notes:** Corresponding Author: Julie K. Andersen.

## Abstract

**Background:** Alzheimer’s disease (AD) is classically defined by amyloid and tau pathology and is accompanied by broad disruptions in proteostasis. Heat shock proteins (HSPs) help maintain proteostasis, yet mitochondrial chaperone systems remain comparatively underexplored in AD. Hsp60 and Hsp10 form a mitochondrial chaperonin complex that folds dozens of AD-implicated mitochondrial proteins, but this client network has not been evaluated as an integrated proteostasis axis in AD. It remains unknown whether Hsp60/10 client proteins are selectively vulnerable across AD severity.

**Methods:** We analyzed transcriptomic, proteomic, neuropathological, and cognitive data from the Religious Order Study and Memory and Aging Project (ROSMAP) to evaluate Hsp60/10 client proteins in AD. We compared Hsp60/10 clients with abundance-matched non-client mitochondrial proteins and tested differences across AD diagnostic groups and associations with Braak/tau burden, cognitive outcomes, and network centrality. These evidence layers were integrated into a candidate prioritization framework.

**Results:** Hsp60/10 client abundance declined more strongly at the protein level than at the RNA level in late-stage AD. Compared with abundance-matched non-client mitochondrial proteins, Hsp60/10 clients showed stronger late-stage protein abundance decline. Greater late-stage client decline was associated with higher Hsp60/10 network centrality, defining a selectively vulnerable client subnetwork. Lower client abundance was associated with greater Braak/tau burden and greater cognitive impairment. Integrated prioritization nominated mitochondrial translation and TCA/pyruvate/redox clients as high-priority candidates for mechanistic follow-up.

**Conclusions:** Together, these findings identify an Hsp60/10 client-centered mitochondrial proteostasis axis spanning mitochondrial translation and TCA/pyruvate/redox metabolism that is associated with AD severity. These findings identify a novel potential axis warranting further investigation as a mechanistic link between mitochondrial dysfunction, proteostasis, and AD.

## INTRODUCTION

Alzheimer’s disease (AD) is a progressive neurodegenerative disorder that affects millions worldwide, imposing substantial emotional and economic burden on patients, families, and caregivers. Its prevalence is expected to rise sharply in the coming decades [1]. Despite extensive study, key gaps remain in our understanding of the molecular mechanisms driving disease pathology.

AD pathology is classically defined by the accumulation of amyloid-β plaques and neurofibrillary tangles, reflecting fundamental disruptions in protein aggregation and cellular proteostasis [2]. Because they promote proper protein folding, heat shock proteins (HSPs) play a central role in maintaining proteostasis. The majority of studies to date have focused on the potential role of cytosolic chaperones such as Hsp70 and Hsp90, while mitochondrial chaperone systems remain comparatively underexplored as potential therapeutic targets in AD [3]. This gap is notable given that mitochondrial dysfunction is a core feature of AD and is linked to impaired protein quality control within the organelle. The Hsp60/10 mitochondrial chaperonin complex, composed of Hsp60 and Hsp10, is responsible for folding more than 300 mitochondrial matrix proteins [4], positioning it as a key regulator of mitochondrial proteostasis.

Hsp60 and Hsp10 have been individually implicated in AD-relevant processes, including mitochondrial dysfunction, proteostasis imbalance, and associated pathology [5,6]. Beyond their canonical role as components of the mitochondrial chaperonin complex, both proteins have also been demonstrated to participate in chaperonin-independent stress-response pathways, including mitochondrial stress signaling functions [7,8]. Several Hsp60/10 client proteins have additionally been independently linked to AD. Despite these connections, prior studies have largely focused on Hsp60, Hsp10, or their client proteins in isolation. The Hsp60/10 chaperonin complex instead represents a unified proteostasis module that functionally connects dozens of AD-implicated mitochondrial proteins. Because Hsp60/10 functions after translation to support mitochondrial protein folding, client protein abundance may provide a more direct readout of chaperonin-related vulnerability than transcript abundance alone. If protein-level loss within this client network is disease-relevant, it would be expected to be associated not only with the AD diagnostic stage, but also with neuropathological burden and cognitive impairment. This client network has not been directly evaluated as an integrated mitochondrial proteostasis axis in AD, representing a critical gap in our understanding of how mitochondrial proteostasis contributes to disease progression.

To address this, we analyzed transcriptomic, proteomic, neuropathological, and cognitive data from the Religious Order Study and Memory and Aging Project (ROSMAP). We tested whether Hsp60/10 client proteins show selective protein-level vulnerability in AD compared to abundance-matched non-client mitochondrial proteins, and whether this vulnerability is associated with neuropathological and cognitive severity. We subsequently prioritized specific client proteins by integrating multiple lines of evidence including network centrality and target nomination status from Agora. This approach enabled the development of a candidate prioritization framework to identify high-value targets for follow-up, providing a strong rationale for future experimental studies to investigate the mechanistic contribution of prioritized Hsp60/10 clients to AD-relevant mitochondrial proteostasis failure.

## METHODS

### Study design and analytical overview

We performed a computational multiomic analysis of bulk RNA-seq and Tandem Mass Tag (TMT) proteomics data to evaluate whether Alzheimer’s disease progression is associated with selective vulnerability of the mitochondrial Hsp60/10 client protein network. Primary discovery analyses were conducted in the Religious Orders Study and Memory and Aging Project (ROSMAP) cohort. A published inventory of mitochondrial Hsp60/10 client proteins was integrated with ROSMAP transcriptomic, proteomic, clinical diagnostic, neuropathologic, and cognitive data (**Fig. S1B**).

The workflow was designed to distinguish broad mitochondrial remodeling from Hsp60/10 client-specific vulnerability (**Fig. S1A**). Bulk RNA-seq provided tissue-level transcript abundance measures aligned with cohort-level proteomics, diagnostic, neuropathologic, and cognitive metadata. Analyses first compared RNA– and protein-level pathway remodeling across clinically defined Alzheimer’s disease stages. ROSMAP-detected Hsp60/10 client proteins were then evaluated for late-stage protein collapse, RNA-protein discordance, protein-pathology coupling, protein-cognition associations, and protein-network centrality. Gene-level protein abundance was modeled in relation to Braak/tau and CERAD/amyloid pathology to evaluate whether client vulnerability was preferentially coupled to tau-related neuropathology. Protein-level cognition models tested whether individual client proteins were associated with cognitive outcomes after accounting for demographic, technical, and pathology-related covariates. Abundance-matched mitochondrial null analyses tested whether Hsp60/10 client signals exceeded empirical expectations from non-client mitochondrial proteins with similar abundance properties. Participant-level Hsp60/10 client abundance scores were analyzed in relation to cognitive diagnosis and cognitive performance. Finally, internal vulnerability metrics, cognition support, network context, Agora nomination status, and MitoCarta-derived functional annotations were integrated using a rule-based framework to classify candidate client proteins for mechanistic follow-up.

Supplemental analyses evaluated robustness and contextual support, including matched-individual RNA/protein sensitivity analyses, alternative pathology-model specifications, cross-cohort validation in Mount Sinai Brain Bank (MSBB) proteomics, and AMP-AD Diverse Cohorts regional proteomics validation.

### ROSMAP cohort, –omics datasets, and sample inclusion

Primary discovery analyses used ROSMAP bulk RNA-seq and TMT proteomics data, and clinical diagnostic groups were defined using the ROSMAP cogdx. Participants with cogdx = 1 were classified as NCI, corresponding to no cognitive impairment. Participants with cogdx = 2 were classified as MCI, corresponding to mild cognitive impairment without another identified cause of cognitive impairment. Participants with cogdx = 4 were classified as AD, corresponding to Alzheimer’s dementia without another identified cause of cognitive impairment. Participants with cogdx = 3, 5, or 6 were excluded from primary stage-associated analyses as these categories indicate cognitive impairment linked to another cause or another primary cause of dementia.

Analyses were restricted to participants with cognitive diagnosis variable (cogdx) values of 1, 2, or 4 and required modality-specific omics data and covariates. For bulk RNA-seq analyses, the primary stage-group sample sizes were 200 NCI, 158 MCI, and 220 AD samples. For TMT proteomics analyses, the primary stage-group sample sizes were 71 NCI, 112 MCI, and 96 AD samples (**Fig. S1C**). These modality-specific sample sets were not restricted to individuals with both RNA-seq and proteomics data, allowing each modality to use the largest available ROSMAP sample set after diagnostic and covariate filtering.

A matched-individual RNA/protein subset was analyzed separately as a sensitivity analysis restricted to participants with both bulk RNA-seq and TMT proteomics data. MSBB proteomics and AMP-AD Diverse Cohorts regional proteomics summary tables were used only after the primary ROSMAP analyses for external validation and contextual support; they were not used to define the primary ROSMAP discovery metrics.

### Neuropathology variables and covariates

Neuropathology analyses focused on Braak stage and CERAD neuritic plaque scores. Braak stage was used as the primary tau-related pathology variable and CERAD score as the primary amyloid-related pathology variable. Braak/tau and CERAD/amyloid associations were modeled separately in the primary pathology-coupling analyses. CERAD-adjusted Braak models were used only as robustness analyses to test whether Braak-associated protein effects persisted after accounting for neuritic plaque burden.

Covariate adjustment was analysis-specific (**Fig. S1D**). For matrix-level preprocessing, bulk RNA-seq abundance values were residualized for age at death, sex, postmortem interval, and RNA integrity number. TMT proteomics abundance values were residualized for age at death, sex, postmortem interval, and protein batch. Diagnostic stage, Braak stage, and CERAD score were not regressed out during matrix-level residualization; these variables were preserved for downstream stage, pathology, cognitive, and robustness analyses.

### Bulk RNA-seq and TMT proteomics matrix preprocessing

ROSMAP RNA abundance was analyzed from a gene-symbol matrix of DESeq2 variance-stabilizing transformed values. Gene identifiers were cleaned and converted to canonical gene symbols. Duplicate rows mapping to the same canonical gene symbol were collapsed by averaging abundance values across duplicated rows within each sample.

ROSMAP TMT proteomics abundance was analyzed from a median-polish corrected log2 abundance matrix. Protein abundance columns were matched to proteomics metadata using TMT batch-channel identifiers and renamed to sample identifiers. RNA and protein matrices were aligned to their corresponding metadata before covariate residualization.

Matrix-level covariate residualization was performed using gene-wise or protein-wise linear models for the specified covariates, with residuals added back to each feature’s mean abundance. The resulting covariate-adjusted matrices were used for stage-associated pathway analyses, collapse, discordance, centrality, and related abundance-summary analyses. No imputation was applied. Individual gene-level models were fitted only when the required variables had sufficient complete observations; otherwise, model-derived values were returned as missing.

### Hsp60/10 client inventory and comparator gene-set definitions

The primary Hsp60/10 client network was defined using a published Hsp60/10 client inventory. Gene identifiers were cleaned and converted to canonical gene symbols; missing, invalid, and duplicate entries were removed. HSPD1 and HSPE1 were excluded because they encoded the Hsp60/10 chaperonin components rather than client proteins. After these exclusions, the reference inventory contained 321 Hsp60/10 clients.

Protein detection-rate categories were summarized for the full 321-client Hsp60/10 reference inventory and for 880 non-client mitochondrial proteins before downstream detection filtering. Detection-rate bins were defined as 0%, >0–50%, 50–95%, and ≥95% detection across ROSMAP TMT proteomics samples. These detection-rate summaries were used descriptively to evaluate protein coverage differences between Hsp60/10 clients and the broader non-client mitochondrial background.

Client detection was determined separately by intersecting the 321 canonical client gene symbols with the row names of the corresponding RNA or protein matrix. Of these clients, 282 were detected in both ROSMAP bulk RNA-seq and TMT proteomics, 12 only in bulk RNA-seq, 21 only in proteomics, and 6 in neither modality. Main all-client RNA analyses therefore used 294 RNA-detected clients, whereas main proteomics analyses used 306 protein-detected clients.

Comparator gene sets were constructed to distinguish Hsp60/10 client-associated effects from broader mitochondrial and proteostasis-related remodeling. Broad mitochondrial comparator genes were derived from Human MitoCarta 3.0 after removing all Hsp60/10 clients. Additional non-client mitochondrial comparator sets were generated from MitoCarta pathway, submitochondrial localization, and gene-description annotations, including mitochondrial translation, oxidative phosphorylation/electron transport chain, TCA/pyruvate metabolism, fatty-acid oxidation/metabolism, and mitochondrial protein quality-control categories. Proteasome and lysosome core comparator sets were manually defined as non-mitochondrial proteostasis references. The mitochondrial unfolded protein response comparator set was manually curated because of overlap with Hsp60/10 clients; client genes were excluded from this set. All pathway-level analyses used only genes or proteins detected in the corresponding matrix.

Detected pathway coverage was recorded separately for RNA-seq and proteomics. The detected RNA/protein feature counts were 294/306 for Hsp60/10 clients, 822/607 for broad mitochondrial non-Hsp60/10 proteins, 121/94 for mitochondrial translation non-Hsp60/10 proteins, 122/94 for OXPHOS/ETC non-Hsp60/10 proteins, 73/66 for TCA/pyruvate non-Hsp60/10 proteins, 95/73 for fatty-acid oxidation/metabolism non-Hsp60/10 proteins, 67/52 for mitochondrial protein quality-control non-Hsp60/10 proteins, 45/45 for proteasome core proteins, 64/45 for lysosome core proteins, and 13/6 for UPRmt core proteins. Full pathway membership and detected coverage are provided in Supplementary Table 1.

### Pathway and participant-level Hsp60/10 client scoring

Pathway abundance scores were calculated separately for bulk RNA-seq and TMT proteomics. For each pathway, detected members were extracted from the corresponding covariate-adjusted matrix and standardized across samples by row-wise z-scoring. Sample-level pathway scores were calculated as the unweighted mean of z-scored abundance values across detected pathway members. This generated mean standardized abundance scores rather than summed or raw abundance measures and prevented features with larger abundance scales or variance from dominating pathway scores. Pathways with fewer than two detected members in a modality were not assigned a score.

The same scoring approach was applied to the Hsp60/10 client network and comparator pathways. RNA and protein scores were calculated within their respective matrices and summarized by diagnostic group using means and standard errors of the mean.

For cognition analyses, a participant-level Hsp60/10 client abundance score was calculated from ROSMAP TMT proteomics data as the mean of standardized abundance values across the 306 ROSMAP-detected client proteins. Higher scores indicate higher relative abundance of the Hsp60/10 client protein network within a participant. Because this score was calculated from z-scored protein abundances, a score of zero represents the cohort-centered client network reference level rather than the expected mean within each diagnostic group.

### Stage-associated RNA and protein remodeling analyses

Stage-associated remodeling analyses were performed separately for bulk RNA-seq and TMT proteomics using the covariate-adjusted matrices. For visualization and comparison, diagnostic labels were harmonized to NCI, MCI, and AD. In bulk RNA-seq analyses, these groups corresponded to cogdx values of 1, 2, and 4, respectively; cogdx = 2 was labeled Early AD in some RNA analysis objects but displayed as MCI in the manuscript. In proteomic analyses, NCI, MCI, and AD corresponded to samples labeled Control, AsymAD, and AD in the proteomics diagnostic metadata.

For each gene, protein, or pathway score, mean abundance was calculated within each diagnostic group. Stage-transition effects were defined as MCI minus NCI for the early-stage effect, AD minus MCI for the late-stage effect, and AD minus NCI for the total AD-associated effect. These effects were calculated independently for RNA and protein features. Protein-minus-RNA remodeling values were calculated for matched pathways to compare the magnitude of stage-associated remodeling between modalities. These stage-transition analyses were kept analytically distinct from gene-level neuropathology-coupling models.

### Late-stage protein collapse, RNA-protein discordance, and network centrality

Late-stage protein collapse was defined as the magnitude of protein abundance decline from MCI to AD. For each ROSMAP-detected protein, the late-stage protein effect was calculated as mean covariate-adjusted protein abundance in AD samples minus mean abundance in MCI samples. Collapse magnitude was calculated as the absolute value of this effect only for proteins with lower mean abundance in AD than MCI; non-declining proteins were assigned a collapse value of zero. Hsp60/10 clients in the top quartile of collapse magnitude were labeled top-quartile collapsing clients for descriptive stratified analyses; this threshold was not treated as a discrete biological boundary.

Late RNA-protein discordance was calculated for Hsp60/10 clients detected in both modalities as the absolute difference between the late-stage protein effect and the corresponding late-stage bulk RNA effect. Higher values indicate a larger mismatch between protein-level and transcript-level remodeling across the MCI-to-AD transition. Discordance was used as a descriptive cross-modality metric and was not interpreted as direct evidence of post-transcriptional regulation, altered protein turnover, or translational control without experimental validation.

Hsp60/10 client network centrality was calculated from the ROSMAP covariate-adjusted TMT proteomics matrix. Pairwise Spearman correlations were computed across proteomic samples for all detected Hsp60/10 client proteins using pairwise complete observations, with self-correlations excluded. For each client protein, centrality was defined as the mean absolute Spearman correlation between that protein and all other detected client proteins. Higher centrality indicates stronger abundance covariation with the broader detected client network.

### Gene-level Braak/tau and CERAD/amyloid pathology coupling

Gene-level protein-pathology coupling analyses were performed across ROSMAP-detected Hsp60/10 client proteins using the processed TMT proteomic abundance matrix and protein covariates. For each detected client protein, abundance values were z-scored across samples before pathology modeling. Braak/tau coupling was estimated from a gene-level linear model relating standardized protein abundance to standardized Braak stage, with age at death, sex, postmortem interval, and protein batch included as covariates. CERAD/amyloid coupling was estimated separately using standardized CERAD neuritic plaque score with the same covariate set.

Pathology effects were direction-oriented so that larger positive values indicate stronger inverse pathology coupling. Negative Braak coefficients, corresponding to lower protein abundance with higher Braak stage, were converted to positive inverse Braak beta values; non-negative coefficients were assigned an inverse Braak beta of zero. CERAD effects were oriented analogously. Braak-minus-CERAD pathology-coupling bias was calculated as inverse Braak/tau coupling minus inverse CERAD/amyloid coupling for each client protein. Positive values indicate stronger inverse coupling with Braak/tau pathology than with CERAD/amyloid pathology. Spearman-based inverse Braak metrics and CERAD-adjusted Braak models were used only in robustness and validation analyses and were not treated as interchangeable with the primary regression-based pathology-coupling estimates.

### Abundance-matched mitochondrial null testing

Abundance-matched mitochondrial null testing was used to determine whether ROSMAP-detected Hsp60/10 client proteins showed stronger vulnerability– and cognition-related signals than non-client mitochondrial proteins with comparable abundance properties. In the final ROSMAP proteomics null framework, the observed set contained 306 detected Hsp60/10 clients, and the matched background pool contained 609 detected non-client mitochondrial proteins from Human MitoCarta 3.0 after removal of Hsp60/10 clients.

For each protein in the null framework, a matching abundance value was calculated from protein abundance across NCI, MCI, and AD groups, and proteins were assigned to abundance bins. Empirical null sets were generated by repeatedly sampling non-client mitochondrial proteins from corresponding abundance bins to match the size and abundance-bin structure of the observed Hsp60/10 client set. Null-set means were calculated for late-stage protein collapse, inverse Braak/tau association, cognition support, and Agora target fraction.

Observed-to-null ratios were calculated as the observed Hsp60/10 client mean divided by the mean of the corresponding matched-null distribution. Empirical p-values were calculated from 10,000 abundance-matched null iterations as the fraction of null sets with values at least as large as the observed Hsp60/10 client value, bounded by the number of iterations.

Cognition support replaced the prior pathology vulnerability score in the abundance-matched null framework. Protein-level cognition support summarized oriented evidence that higher client protein abundance was associated with better cognitive outcomes across the cognition models described below. This metric was included to test whether Hsp60/10 clients were enriched for proteins linked to cognitive preservation, rather than only for proteins showing combined late-stage decline and Braak-associated decline. For the null framework, the cognition preservation score was calculated from the summarized oriented cognition-support ranking across the three protein-level cognition models, with higher values indicating stronger association between higher client abundance and better cognitive status.

### Cognitive outcome and protein-level cognition-support analyses

Cognitive outcome analyses tested whether Hsp60/10 client protein abundance was associated with cognitive diagnosis and cognitive performance in ROSMAP. Participant-level analyses used an Hsp60/10 client abundance score calculated as the mean of standardized abundance values across the 306 ROSMAP-detected client proteins. Each participant’s client abundance score was analyzed in relation to three cognition-related outcomes: diagnosis at death, last-valid cognitive diagnosis, and last-valid Mini-Mental State Examination score. Cognitive preservation score was defined as the relationship between higher client abundance and better cognition/lower impairment, oriented so positive values are associated with better cognition. Diagnosis-at-death groups were also visualized as no cognitive impairment, mild cognitive impairment, and Alzheimer’s disease dementia.

Associations between participant-level Hsp60/10 client abundance and cognitive outcomes were estimated using regression models that included age at death, sex, education, postmortem interval, Braak stage, and CERAD neuritic plaque score as covariates. For visualization of diagnosis-at-death groups, the Hsp60/10 client score was adjusted for these covariates and plotted across diagnostic categories. Participants with complete proteomics, diagnosis, pathology, and covariate data were grouped as no cognitive impairment, NCI (n = 103); mild cognitive impairment, MCI (n = 62); or Alzheimer’s disease dementia, AD (n = 66). For visualization of the association between Hsp60/10 client abundance and last-valid Mini-Mental State Examination score, both variables were residualized with respect to the covariates and pathology variables included in the model; these partial residuals were plotted against each other to display the adjusted association.

Protein-level cognition-support analyses were performed across ROSMAP-detected Hsp60/10 client proteins using the processed TMT proteomics abundance matrix. For each protein, standardized abundance was modeled in relation to diagnosis at death, last-valid cognitive diagnosis, and last-valid Mini-Mental State Examination score, with higher outcome values corresponding to better cognitive status. Models included age at death, sex, education, postmortem interval, Braak stage, and CERAD neuritic plaque score as covariates. Protein-level cognition effects were oriented so that larger positive values indicated stronger association between higher protein abundance and better cognition. Cognition-support evidence was summarized across the three cognitive outcomes and converted to within-client percentile ranks for abundance-matched null testing and integrated candidate classification.

For visualization of cognition-prioritized clients, proteins were first filtered for FDR-supported associations across all three cognition-related outcomes and then ranked by the summarized cognition-support metric. The top 25 ranked clients were displayed in **Fig. 4B**. For class-level summaries, each detected client was assigned to a MitoCarta functional class and categorized by the number of cognition outcomes reaching FDR support: 3/3, 2/3, 1/3, or 0/3 outcomes.

### Integrated candidate classification and rule-based prioritization

Integrated candidate classification was performed to organize ROSMAP-detected Hsp60/10 client proteins for mechanistic follow-up. The primary internal evidence axes were late-stage protein collapse, inverse Braak/tau coupling, cognition support, and Hsp60/10 client network centrality. Each axis was converted to a within-client percentile rank so that measures with different raw units could be compared on a common scale. For an external support annotation, we used Agora-nominated target status. Agora hosts high-dimensional human transcriptomic, proteomic, and metabolomic evidence for whether genes are associated with AD. It contains a list of more than 900 nascent AD drug targets nominated by researchers from the National Institute on Aging’s AMP-AD consortium and TREAT-AD centers, as well as by other AD research teams. Agora status was not used as a prerequisite for candidate classification. MitoCarta 3.0-derived annotations provided functional context.

Pareto/frontier status was calculated from collapse percentile, inverse Braak/tau percentile, cognition percentile, and centrality percentile. Pareto/frontier candidates were defined as proteins with non-dominated combinations of these four axes, meaning no other detected client showed equal or stronger support across all four axes and stronger support on at least one axis. Agora support was then used to subdivide candidate layers into internally prioritized clients with Agora support, internally prioritized clients without Agora support, Agora-supported non-frontier clients, and other classified clients. Candidate classes were intended to organize proteins for downstream experimental evaluation, not to establish causality or therapeutic efficacy.

For functional-class summaries, MitoCarta classes were summarized by mean collapse percentile, mean inverse Braak/tau percentile, mean cognition percentile, mean centrality percentile, Pareto/frontier fraction, and Agora fraction. Candidate-layer display tables showed up to six representative genes per layer. Within each layer, rows were ordered for visualization using the unweighted mean of collapse, inverse Braak/tau, cognition, and centrality percentiles; this display ordering was not used to define Pareto/frontier status or candidate-layer assignment. The full ranked candidate table is provided in Supplemental Table 5.

### Matched-individual bulk RNA/proteomics sensitivity analysis

A matched-individual sensitivity analysis tested whether RNA/protein remodeling patterns were preserved when analyses were restricted to ROSMAP participants with both bulk RNA-seq and TMT proteomics data. The primary stage-associated remodeling analyses used the largest available modality-specific sample sets, whereas this sensitivity analysis used only individuals represented in both modalities. Hsp60/10 client and comparator pathway scores were calculated using the same mean z-scored abundance approach described above, separately within each modality, and compared across harmonized NCI, MCI, and AD groups.

For the matched-individual sensitivity analysis, sample availability varied by modality and pathway after requiring matched omics and detected pathway members. The Hsp60/10 client network included RNA samples from NCI (n = 65), MCI (n = 41), and AD (n = 35) groups and protein samples from NCI (n = 45), MCI (n = 59), and AD (n = 38) groups. The broad mitochondrial background included RNA samples from NCI (n = 65), MCI (n = 41), and AD (n = 36) groups and protein samples from NCI (n = 45), MCI (n = 59), and AD (n = 38) groups. The TCA/pyruvate non-Hsp60/10 comparator included RNA samples from NCI (n = 54), MCI (n = 41), and AD (n = 36) groups and protein samples from NCI (n = 45), MCI (n = 59), and AD (n = 38) groups.

### Pathology-model robustness analyses

Pathology-model robustness analyses evaluated whether Hsp60/10 client protein-pathology coupling was preserved across alternative Braak/tau association metrics and model specifications. Unadjusted Spearman correlations were used to estimate monotonic associations between protein abundance and Braak stage, with effects direction-oriented so that larger positive values indicated stronger inverse association. Covariate-adjusted Braak models estimated inverse Braak/tau coupling using standardized protein abundance and standardized Braak stage with age at death, sex, postmortem interval, and protein batch as covariates. CERAD-adjusted Braak models additionally included CERAD neuritic plaque score to test whether Braak-associated protein effects persisted after accounting for neuritic plaque burden.

Robustness analyses compared ROSMAP-detected Hsp60/10 clients with detected non-client mitochondrial proteins across Spearman, covariate-adjusted, and CERAD-adjusted Braak specifications. Late-stage protein collapse was also visualized against adjusted inverse Braak/tau coupling to evaluate whether stage-associated decline and pathology-associated decline identified overlapping client proteins.

### External MSBB cross-cohort validation

External cross-cohort validation was performed using MSBB proteomics data to evaluate whether Hsp60/10 client pathology-coupling patterns observed in ROSMAP were directionally reproduced in an independent cohort. MSBB analyses used a normalized TMT proteomics abundance matrix provided as a gene-level spreadsheet and were treated as validation and contextual support, not as part of the primary ROSMAP discovery framework.

Gene identifiers were harmonized to canonical gene symbols. Hsp60/10 clients and non-client mitochondrial proteins were defined using the same client inventory and mitochondrial-background logic applied in ROSMAP, with analyses restricted to proteins detected in the MSBB proteomics matrix. MSBB protein-pathology effects were oriented so that larger positive values indicated stronger inverse association between protein abundance and Braak/tau pathology. Cross-cohort concordance was defined as inverse Braak/tau association in both ROSMAP and MSBB for the same protein.

For gene-level cross-cohort concordance analyses, Hsp60/10 clients were restricted to proteins detected in both ROSMAP and MSBB proteomics. Concordance categories were defined according to whether each detected client showed inverse Braak/tau association in both cohorts, ROSMAP only, MSBB only, or neither/opposite direction. The concordance visualization included 290 Hsp60/10 clients detected in both cohorts.

### Regional proteomics validation

Regional proteomics validation used precomputed AMP-AD Diverse Cohorts regional TMT proteomics summary tables from the dorsolateral prefrontal cortex and superior temporal gyrus. These tables contained gene-level AD-associated protein effects, Braak-associated protein effects, and STG-DLPFC regional contrast metrics. Regional analyses were treated as validation and contextual support rather than as part of the primary ROSMAP discovery framework.

Gene identifiers were harmonized to canonical gene symbols. Hsp60/10 clients and non-client mitochondrial proteins were defined using the same client inventory and mitochondrial-background logic used in the primary ROSMAP analyses, with analyses restricted to proteins represented in the regional summary tables. Regional protein effects were oriented so that larger positive values indicated stronger AD-associated protein decline or stronger inverse association between protein abundance and Braak/tau pathology. Analyses compared Hsp60/10 clients with non-client mitochondrial proteins for AD-associated decline, Braak-associated decline, and STG-DLPFC regional differences.

For STG-DLPFC regional contrast analyses, region differences were calculated as the STG-oriented effect minus the DLPFC-oriented effect for each Hsp60/10 client. Positive STG-DLPFC values indicate stronger AD-associated decline or stronger inverse Braak/tau coupling in STG than DLPFC, whereas negative values indicate stronger signal in DLPFC. Hsp60/10 clients displayed in the regional lollipop plot were ranked by STG-DLPFC adjusted inverse Braak magnitude.

### Sensitivity analyses for stage-residualized co-movement and functional specificity

We performed two sensitivity analyses to test whether Hsp60/10 client vulnerability could be explained by shared diagnostic-stage effects or by broad mitochondrial functional class. First, using the covariate-adjusted ROSMAP proteomics matrix, each detected Hsp60/10 client protein was additionally residualized for diagnostic group. Pairwise correlations were then recalculated across residualized client abundance profiles. Mean absolute pairwise residual correlation was calculated for two selected client subsets: the top late-collapsing clients and the collapse-central clients. For each subset, the observed mean absolute residual correlation was compared with size-matched random Hsp60/10 client sets using 1,000 permutations, with random sets matched on available client attributes where possible (e.g. MitoCarta 3.0 functional class). Empirical one-sided p values were calculated as the proportion of matched null sets with equal or greater mean residual co-movement than the observed subset. Across all detected clients, the relationship between late-stage protein-collapse magnitude and mean residual co-movement to other clients was tested using Spearman correlation.

Second, we tested whether client vulnerability exceeded that of non-client mitochondrial proteins with similar functional annotations and abundance. Detected Hsp60/10 clients and detected non-client MitoCarta proteins were assigned to mitochondrial functional modules. Analyses were restricted to modules containing both clients and non-client mitochondrial proteins. For each metric, including late-stage protein-collapse magnitude, inverse Braak association magnitude, and integrated pathology vulnerability score, the observed client mean was compared with function– and abundance-matched non-client mitochondrial null sets using 1,000 one-sided permutations. Module-level analyses were performed for late-stage protein-collapse magnitude. As an additional sensitivity analysis, linear models tested the association between client status and each vulnerability metric while adjusting for functional module and protein abundance. Results from these analyses are reported in **Supplementary Table 7**.

### Statistical testing conventions

Statistical analyses were performed at the pathway, gene/protein, or participant level according to the analysis objective. Unless otherwise specified, two-sided tests were used. Group comparisons between Hsp60/10 clients and non-client mitochondrial proteins used Wilcoxon rank-sum tests. Paired comparisons, including inverse Braak/tau versus inverse CERAD/amyloid coupling within the same proteins, used paired Wilcoxon signed-rank tests.

Regression-based analyses were used when covariate adjustment was required. Gene-level pathology models used linear regression, and participant-level cognitive analyses used regression models appropriate to the outcome. For visualization of adjusted relationships, residualized or partial-residual values were used only for plotting; statistical inference was based on the corresponding regression models.

For gene-level model families, p-values were adjusted using the Benjamini-Hochberg false discovery rate procedure. Abundance-matched mitochondrial null analyses used empirical p-values from matched-null distributions. Direction-oriented vulnerability metrics were constructed so that larger positive values consistently indicated stronger protein decline, stronger inverse pathology association, or stronger vulnerability-related evidence; effects in the opposite direction were assigned values of zero rather than negative vulnerability scores. Nominal significance was defined using conventional thresholds of P < 0.05, P < 0.01, and P < 0.001 where displayed in figures. Statistical results were interpreted alongside effect direction, magnitude, cohort context, and consistency across complementary analyses.

### Software and reproducibility

All analyses were performed using scripted workflows in R. The pipeline was organized into modular scripts for data loading, matrix preprocessing, pathway scoring, stage-associated remodeling, pathology modeling, matched-null testing, cognitive analyses, validation analyses, and figure generation. Shared helper functions were used for gene-symbol cleaning, matrix alignment, covariate residualization, pathway-score calculation, model fitting, and output writing.

Intermediate analysis objects, figure source tables, audit tables, and statistical summaries were written to disk to verify sample inclusion, detection, covariate adjustment, pathway coverage, model completeness, and matched-null construction. Figures were generated from saved analysis tables rather than manually assembled from plotted values. Random sampling procedures used fixed seeds where applicable, including abundance-matched mitochondrial null testing. Multiple-testing correction, empirical p-value calculation, and percentile-rank transformations were performed programmatically. Software dependencies included tidyverse packages for data manipulation and plotting, with additional packages for model tidying, spreadsheet import, figure assembly, and label placement. Custom R analysis code used to generate the manuscript analyses and figures is available at https://github.com/asloane-buck/hsp60-10_rosmap_publication_submission and archived on Zenodo at https://doi.org/10.5281/zenodo.20851582. Controlled-access ROSMAP transcriptomic, proteomic, clinical, neuropathological, and metadata files are available through the applicable AD Knowledge Portal/Synapse access procedures and are not redistributed with the code repository.

## RESULTS

### Stage-associated mitochondrial pathway remodeling is protein-biased but not unique to Hsp60/10 clients

ROSMAP bulk RNA-seq and TMT proteomics were integrated to compare stage-associated RNA and protein abundance changes in various database-derived gene sets across AD progression, defined here as pathway remodeling. We then asked whether pathway remodeling was biased towards protein over RNA. The 306 detected Hsp60/10 client genes displayed statistically significant pathway remodeling for both RNA and protein, as well as RNA-protein discordance in late stages of AD progression. Broad non-client mitochondrial genes showed comparable RNA/protein remodeling and discordance. Therefore, Hsp60/10 clients were not uniquely remodeled relative to broad non-client mitochondrial genes (**Fig. 1A**).

**Figure 1.**
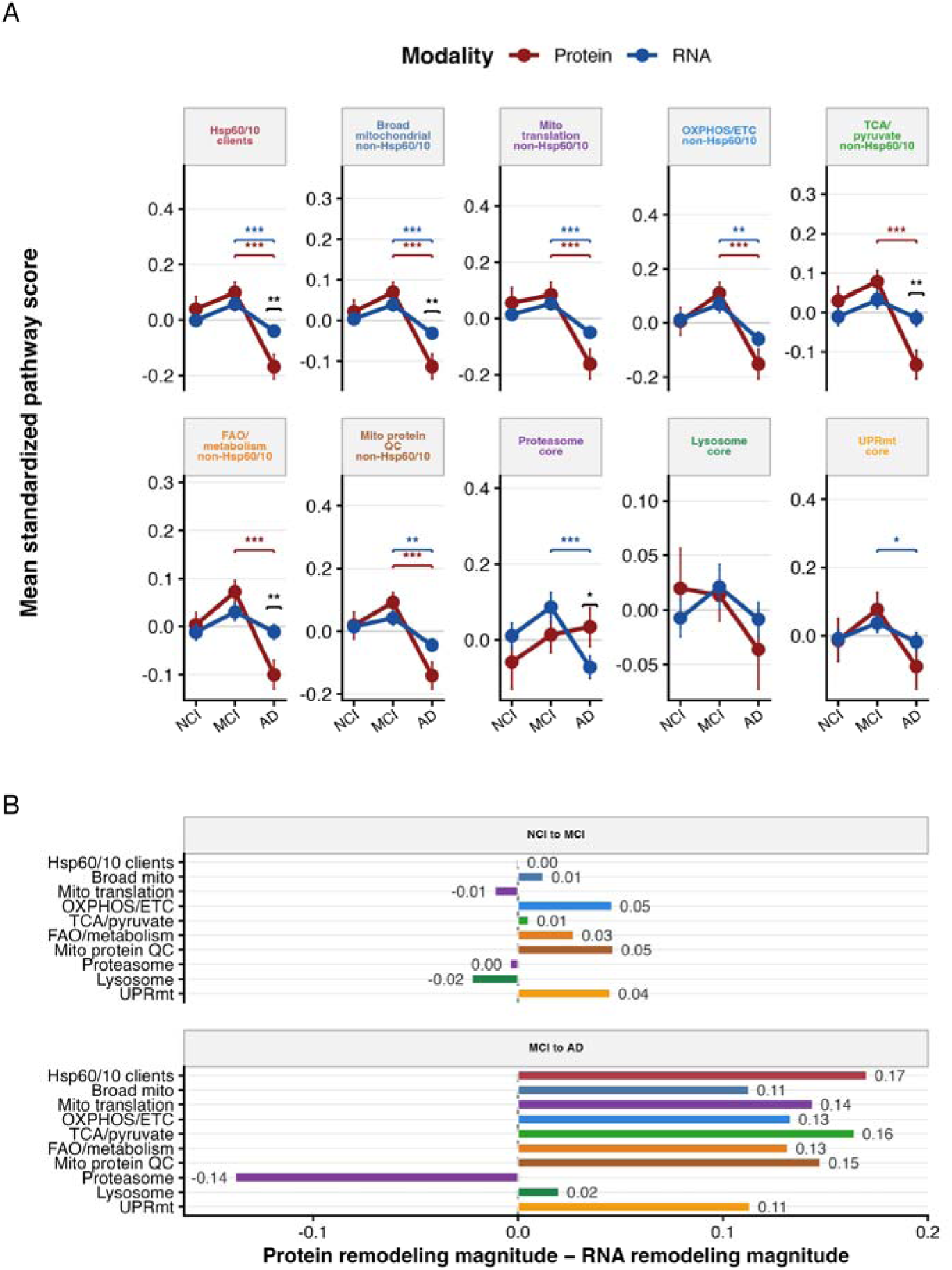
Hsp60/10 analysis resolves protein-biased mitochondrial remodeling. ROSMAP bulk RNA-seq and TMT proteomics were integrated to compare RNA-estimated and protein-level pathway remodeling across NCI, MCI, and AD groups. Sample sizes were n = 200, 158, and 220 for RNA and n = 71, 112, and 96 for protein. Pathway scores represent mean row-standardized abundance across detected pathway members. (A) Mean standardized pathway scores are shown for the detected Hsp60/10 client network and comparator pathways indicated in panel labels. Red indicates protein and blue indicates RNA. Points show group means; error bars show SEM. Colored brackets indicate within-modality stage comparisons; black brackets indicate protein-versus-RNA differences within stage. *P < 0.05, **P < 0.01, ***P < 0.001. (B) Protein-biased remodeling magnitude for the NCI-to-MCI and MCI-to-AD transitions, calculated as protein remodeling magnitude minus RNA remodeling magnitude. Positive values indicate greater stage-related remodeling at the protein level; negative values indicate greater RNA-level remodeling or relative protein-level attenuation. Numeric labels show remodeling differences.

We investigated whether this non-client mitochondrial result was simply an artifact of grouping all mitochondrial proteins together by analyzing more specific database-derived gene sets of non-client mitochondrial genes. Specific comparator sets showed category-dependent remodeling but did not overturn the conclusion that Hsp60/10 clients were not uniquely remodeled relative to non-client mitochondrial proteins.

Matched-individual analyses, used to address incomplete RNA/protein sample overlap, were directionally consistent with the unmatched analyses, which used all available RNA and protein samples (**Fig. S2**).

We then subtracted the magnitude of RNA remodeling from the magnitude of protein remodeling, which revealed that pathway remodeling was biased towards protein, particularly in the MCI-to-AD transition. The degree of protein-biased remodeling was similar across all pathways, with the exception of proteasomal genes (**Fig. 1B**).

Hsp60/10 clients are remodeled, but broad pathway analysis alone does not establish Hsp60/10-specific vulnerability. We next tested whether vulnerability was concentrated among a subset of individual clients.

### Late-stage protein collapse identifies a network-central subset of Hsp60/10 clients

Because pathway-level remodeling did not establish specificity, we tested whether heterogeneity in AD progression was concentrated within individual clients by ranking them by late-stage protein collapse, RNA-protein discordance, and network centrality. Hsp60/10 clients showed heterogeneous late-stage collapse, indicating that the client network does not uniformly decline in protein abundance during the MCI-to-AD transition (**Fig. 2A**). This top-collapsed subset was not distinguished by greater RNA-protein discordance (**Fig. 2B**) but showed higher network centrality, which captures relative connectivity within the Hsp60/10 client network (**Fig. 2C**). Thus, late-stage protein collapse, and this protein-level vulnerability, appears concentrated in a network-central subset of the Hsp60/10 client network.

**Figure 2.**
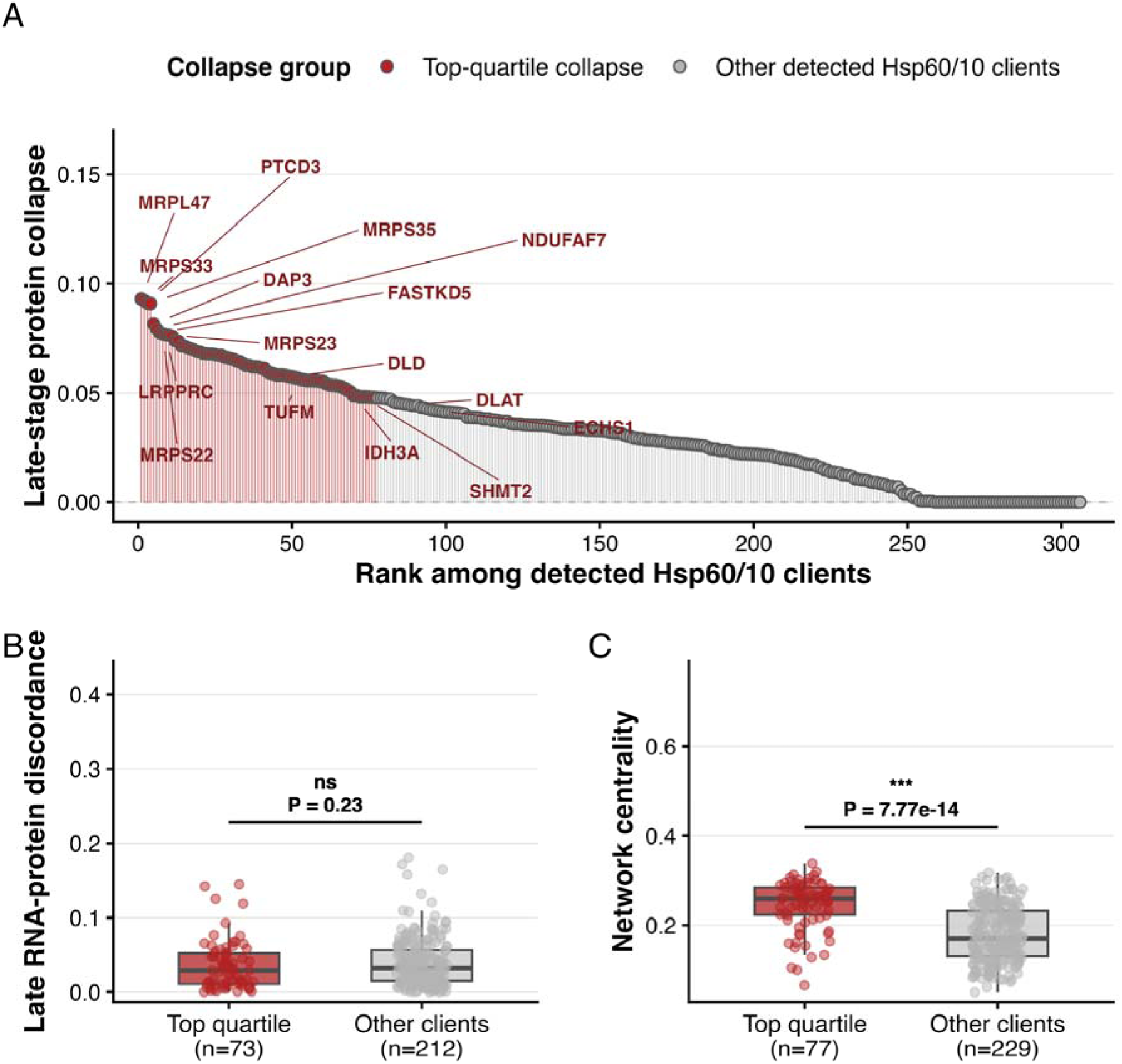
Hsp60/10 client vulnerability is selective across the full client network. Detected Hsp60/10 client proteins in ROSMAP proteomics were ranked by late-stage protein collapse to determine whether vulnerability was broadly distributed or concentrated among strongly declining clients. Collapse was defined as MCI-to-AD protein decline, with non-declining proteins assigned a value of zero. Group comparison statistics are shown in panels. (A) Continuous ranking of late-stage protein collapse across detected clients. Red indicates top-quartile collapsing clients, and gray indicates other detected clients; the quartile threshold is descriptive rather than a biologically discrete boundary. (B) Late RNA-protein discordance compared between top-quartile collapsing clients (n = 73 proteins) and other clients (n = 212 proteins) using a two-sided Wilcoxon rank-sum test, P = 0.23. Discordance reflects mismatch between late-stage protein remodeling and RNA-estimated remodeling; higher values indicate greater RNA-protein mismatch. The smaller n reflects the requirement for matched RNA and protein information. (C) Protein-network centrality compared between top-quartile collapsing clients (n = 77 proteins) and other clients (n = 229 proteins) using a two-sided Wilcoxon rank-sum test, P = 7.77 x 10^−14^. Network centrality captures relative connectivity within the Hsp60/10 client network.

### Hsp60/10 client abundance is more strongly coupled to Braak/tau pathology than CERAD/amyloid pathology

We tested whether Hsp60/10 clients were more strongly coupled to CERAD/amyloid or Braak/tau pathology and whether any pathology correlations were related to late-stage protein collapse. Individual Hsp60/10 client protein abundances were inversely associated with Braak stage (**Fig. 3A**). Higher values indicate stronger association between lower client abundance and greater Braak/tau pathology. The proteins that collapsed most strongly in late-stage AD tended to show stronger inverse Braak/tau coupling (**Fig. 3B)**. Braak/tau coupling exceeded CERAD/amyloid coupling for most clients (**Fig. 3C**).

**Figure 3.**
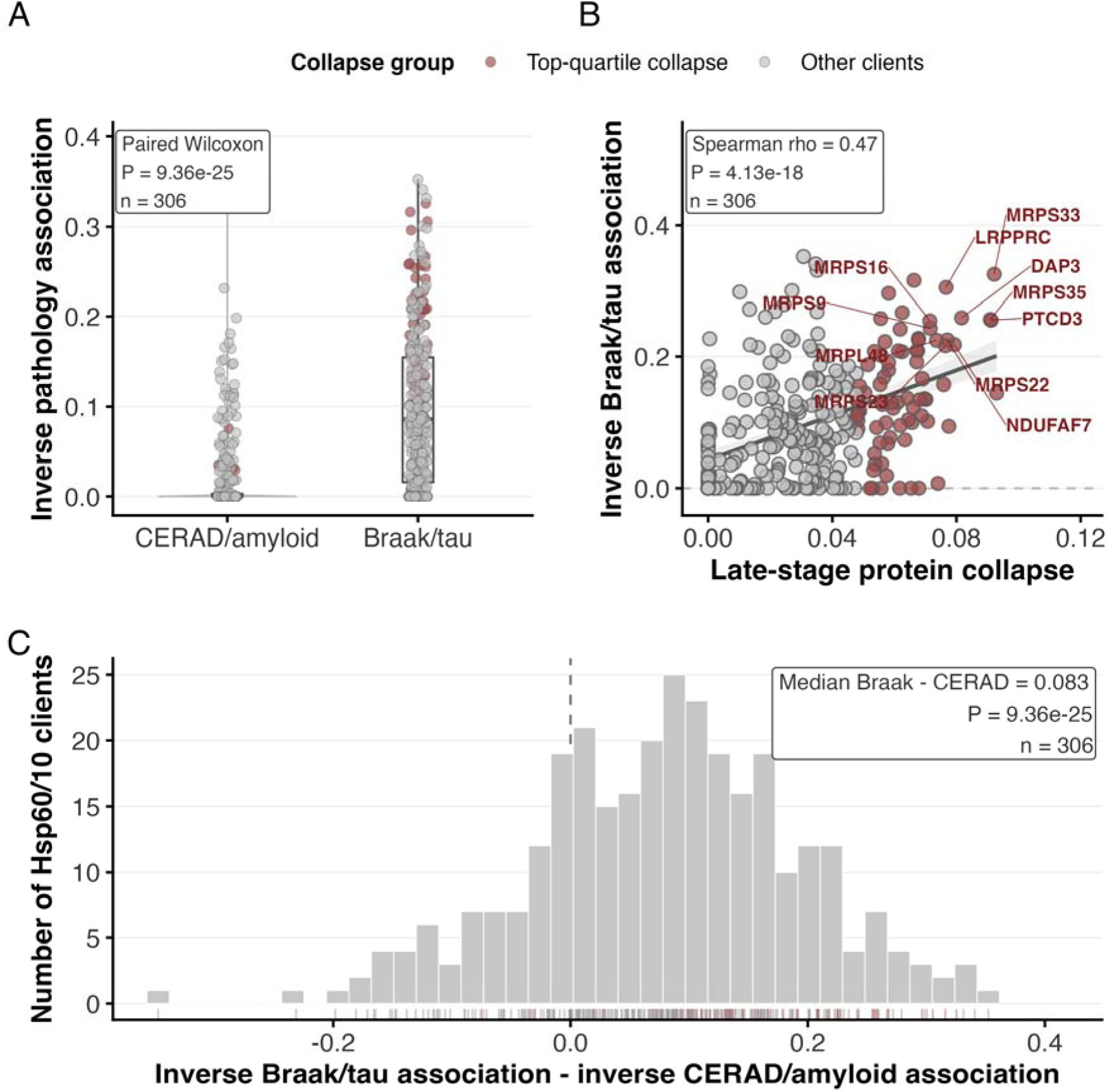
Hsp60/10 client vulnerability is preferentially coupled to Braak/tau pathology. Gene-level pathology associations were calculated across detected Hsp60/10 client proteins to compare coupling with tau-related versus amyloid-related pathology. Protein associations with Braak stage and CERAD score were modeled separately after adjustment for demographic and technical covariates. Associations were direction-oriented, so higher values indicate lower protein abundance with greater pathology burden. Late-stage collapse represents covariate-adjusted MCI-to-AD protein decline. Statistical summaries for paired tests and correlation analyses are shown in panels. (A) Inverse pathology coupling across detected clients (n = 306 proteins) for CERAD/amyloid and Braak/tau using a paired Wilcoxon signed-rank test, P = 9.36 x 10^−25^. Each point represents one client protein; boxplots summarize medians and interquartile ranges. Top-quartile collapsing clients are highlighted in red. (B) Late-stage protein collapse was correlated with inverse Braak/tau coupling across detected clients (n = 306 proteins) using Spearman correlation, rho = 0.47, P = 4.13 x 10^−18^. Red points indicate top-quartile collapsing clients; gray points indicate other detected clients. Labeled proteins identify representative clients with high collapse and/or strong Braak/tau coupling. The fitted line summarizes the continuous association. (C) The within-client Braak-minus-CERAD coupling bias had median difference = 0.083 across n = 306 proteins and was tested using the paired Wilcoxon signed rank test from panel A, P = 9.36 x 10^−25^. The dashed line marks zero; rung marks indicate individual clients, with top-quartile collapsing clients highlighted in red.

Sensitivity models showed that the inverse Braak association pattern remained detectable after conservative covariate adjustment, including models that accounted for CERAD/amyloid burden (**Fig. S4C**).

Independent MSBB proteomics analysis provided modest, directionally consistent support for Hsp60/10 client pathology coupling (**Fig. S5B**). Across the ROSMAP and MSBB cohorts, Hsp60/10 client protein correlations were modestly concordant (**Fig. S5C**). Clients also displayed region-specific patterns, with stronger AD protein collapse and inverse Braak stage association observed in the superior temporal gyrus (STG) than the dorsolateral prefrontal cortex (DLPFC) (**Fig. S6C**).

Overall, the pathology axis supports a stronger relationship between Hsp60/10 client abundance and Braak/tau pathology than CERAD/amyloid pathology, with the strongest collapse-prioritized clients also showing stronger Braak coupling.

### Client abundance tracks cognitive outcomes and prioritizes mitochondrial translation and TCA/pyruvate/redox clients

We next asked whether Hsp60/10 network-level client abundance was associated with cognitive outcomes. Network-level Hsp60/10 client abundance declined with increasing diagnostic severity, as individuals diagnosed with AD at death had lower client abundance than those with no cognitive impairment (NCI) or mild cognitive impairment (MCI) (**Fig. 4A**).

**Figure 4.**
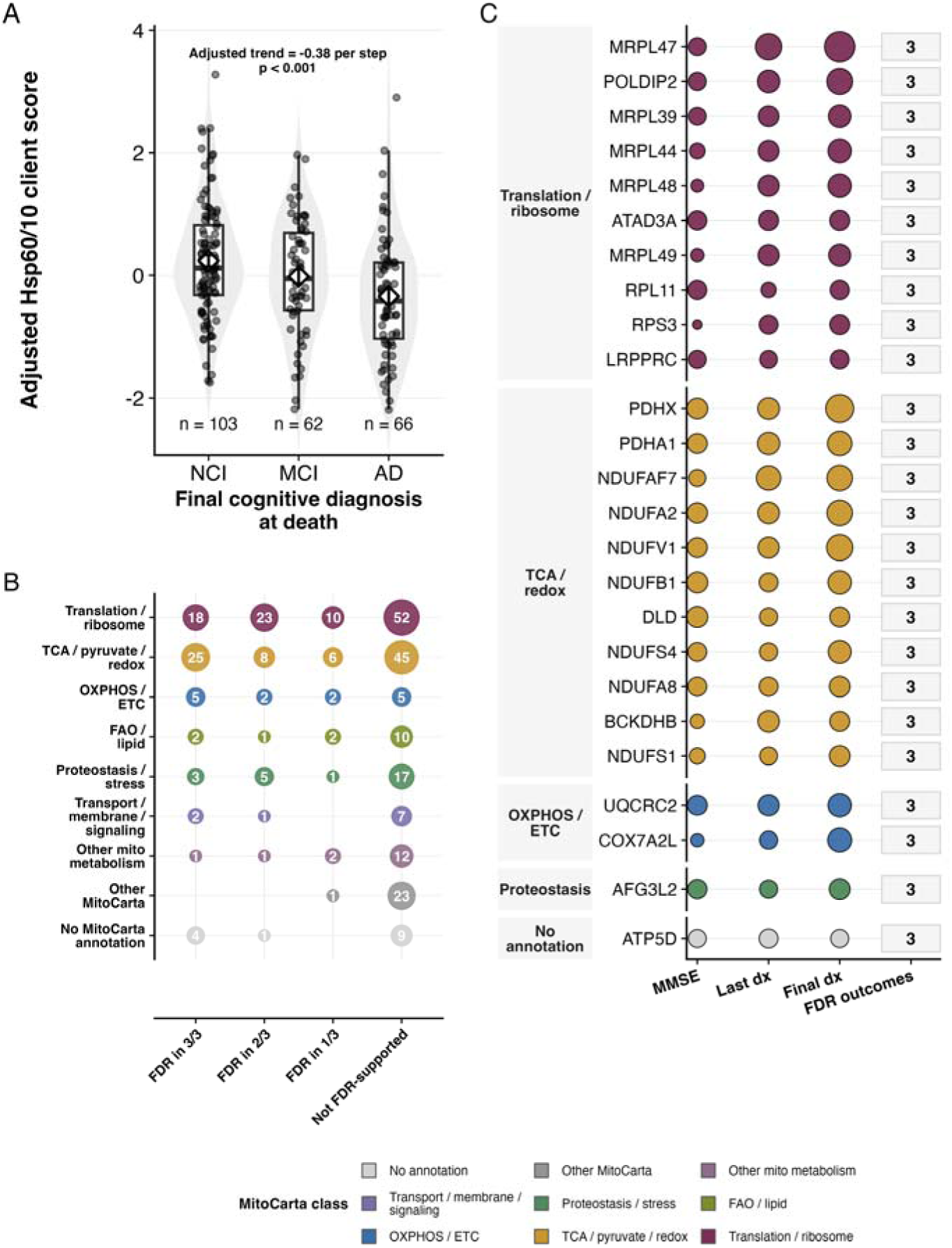
Hsp60/10 client network abundance tracks cognitive preservation. ROSMAP TMT proteomics was used to quantify Hsp60/10 client network abundance across 306 detected client proteins and evaluate associations with cognitive impairment and cognition-related outcomes. Client scores and protein-level models were adjusted for covariates and pathology-related variables. Higher cognitive outcome values indicate better cognitive status. (A) Covariate-adjusted Hsp60/10 client abundance scores across final cognitive diagnosis at death: no cognitive impairment, NCI, n = 103; mild cognitive impairment, MCI, n = 62; and Alzheimer’s disease dementia, AD, n = 66. The participant-level client score was calculated as the mean of z-scored abundance across detected clients. Violin plots show distributions, boxplots summarize medians and interquartile ranges, and points represent participants. Adjusted diagnostic modeling showed decreasing client abundance with worsening cognitive diagnosis, β = −0.38 per diagnostic step, p < 0.001. (B) Top 25 cognition-prioritized clients across last-valid MMSE, last-valid cognitive diagnosis, and final diagnosis at death. Rows represent client proteins grouped by MitoCarta functional class. Dot size represents −log10(FDR), color indicates functional class, and the right column shows the number of FDR-supported outcomes. (C) Detected clients summarized by MitoCarta functional class and number of FDR-supported cognition outcomes. Dot labels indicate the number of detected clients in each class-outcome-support category.

Individual clients were then ranked by their associations across three cognitive outcomes: last-valid MMSE, last-valid cognitive diagnosis, and diagnosis at death. The top 25 cognition-prioritized clients all had FDR-significant associations with all three outcomes (**Fig. 4B**).

Cognition-prioritized clients were concentrated on mitochondrial translation and TCA/pyruvate/redox metabolism functional classes (**Fig. 4C**). Together, these findings identify cognition as an additional axis for client prioritization, extending the client-abundance pattern from neuropathology to cognitive status.

### Abundance-matched mitochondrial null testing supports client specificity across multiple AD-relevant axes

Because prior analyses showed broad mitochondrial remodeling, we next tested whether the client signal exceeded what would be expected from non-client mitochondrial proteins with similar protein abundance. We used mitochondrial null testing to compare Hsp60/10 clients with 10,000 abundance-matched non-client mitochondrial null sets (**Fig. 5A**). Hsp60/10 clients exhibited a 1.25-fold higher late-stage protein collapse and a 1.39-fold higher inverse Braak association. Clients also showed a 1.61-fold higher cognitive preservation score, reflecting stronger association between client abundance and better cognitive status/lower impairment. Clients showed more than a two-fold higher Agora nomination fraction than abundance-matched null sets (**Fig. 5B**). These enrichments indicate that Hsp60/10 clients show group-level specificity across molecular, neuropathological, cognitive, and external target-nomination axes. However, this group-level enrichment did not identify which individual clients should be prioritized.

**Figure 5.**
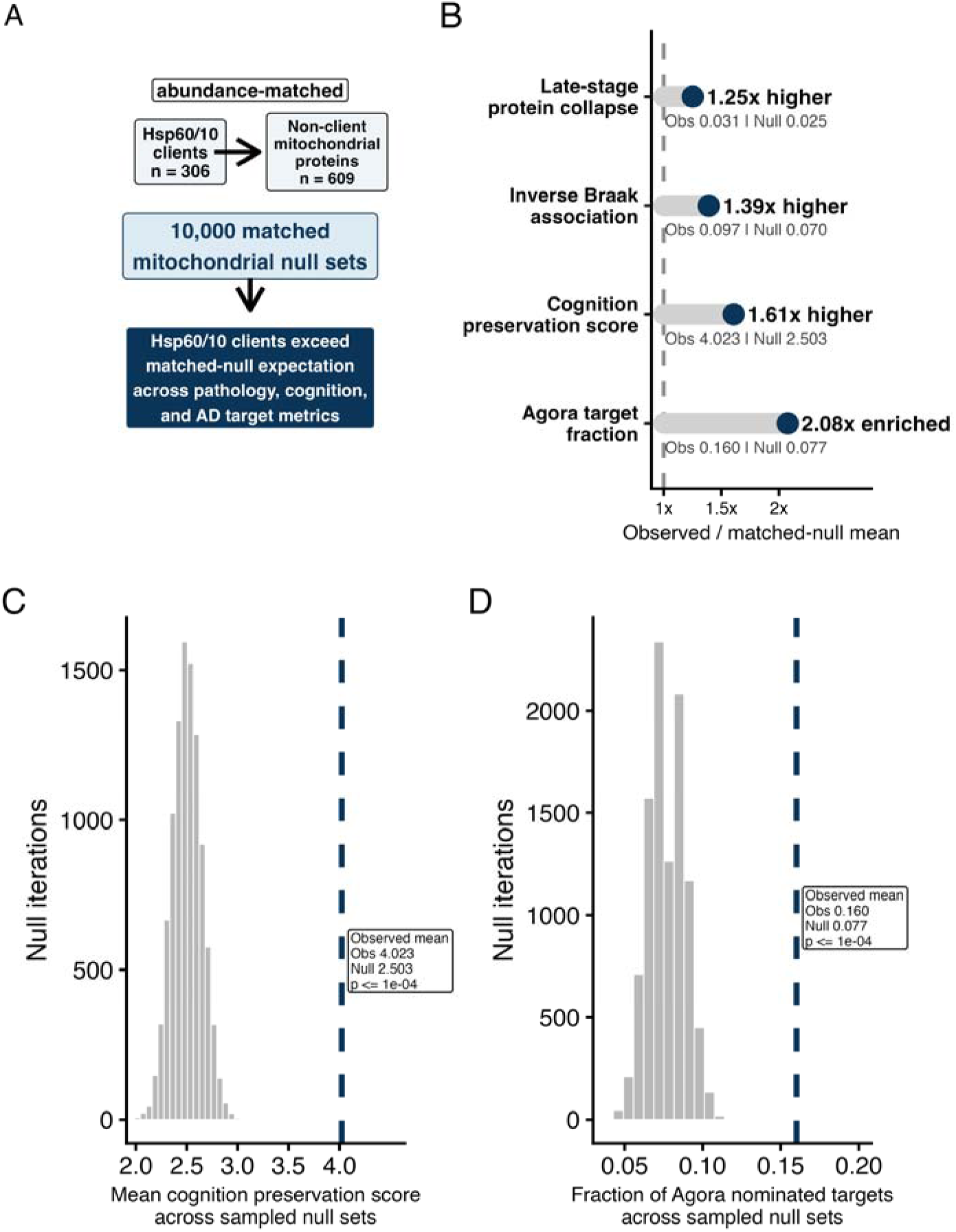
Hsp60/10 clients define a selectively AD– and cognition-relevant mitochondrial subnetwork. A matched-null framework compared the detected Hsp60/10 client network with non-client mitochondrial protein sets of similar abundance. The observed set contained 306 detected clients, and the matched background pool contained 609 detected non-client mitochondrial proteins from MitoCarta 3.0. For null testing, 10,000 abundance-matched mitochondrial null sets were sampled from the non-client background. Observed/null values, enrichment ratios, and empirical p-values are shown in panels. (A) Schematic of the abundance-matched mitochondrial specificity test comparing observed Hsp60/10 client metrics against matched-null distributions. (B) Observed-to-null enrichment ratios were 1.25 for late-stage protein collapse, 1.39 for inverse Braak association, 1.61 for cognition preservation score, and 2.08 for Agora target fraction; empirical P ≤ 1.0 x 10^−4^, with values bound by 10,000 null iterations. Values represent the observed Hsp60/10 client mean divided by the matched mitochondrial null mean. The dashed line marks 1.0, indicating no enrichment. (C) Matched-null distribution for cognition preservation score: observed mean = 4.023, null mean = 2.503, empirical P ≤ 1.0 x 10^−4^, with values bound by 10,000 null iterations. The dashed line marks the observed Hsp60/10 client mean. (D) Matched-null distribution for Agora target fraction: observed fraction = 0.160, null mean = 0.77, empirical P ≤ 1.0 x 10^−4^, with values bound by 10,000 null iterations. The dashed line marks the observed Hsp60/10 client fraction.

### Sensitivity analyses support client vulnerability beyond disease stage and mitochondrial functional class

Additional sensitivity analyses supported the specificity of the client vulnerability signal. After protein abundance was residualized for disease stage, both top late-stage collapsing clients and network-central clients retained higher co-movement than matched client null sets, indicating that the coordinated client signal was not explained solely by diagnostic-stage differences. Across all detected clients, stronger late-stage protein collapse was also associated with greater stage-residualized co-movement. In a separate function-matched analysis, Hsp60/10 clients remained more vulnerable than non-client mitochondrial proteins matched by functional module and abundance, including for late-stage protein collapse and inverse Braak association magnitude. Module-level effects were strongest for mitochondrial translation and TCA/pyruvate metabolism (**Supplementary Table 7**).

### Integrated evidence prioritizes mitochondrial translation and TCA/pyruvate/redox Hsp60/10 clients for follow-up

We combined the preceding client-level evidence axes to create a candidate classification framework to prioritize Hsp60/10 clients for mechanistic follow-up (**Fig. 6A**). The framework incorporated four prioritization evidence axes: late-stage protein collapse (protein vulnerability), inverse Braak/tau association (pathology orientation), client-level cognition association (cognitive preservation), and network centrality. These axes were used to identify Pareto/frontier candidates. Pareto/frontier candidates were defined as clients that were not outperformed by any other client across all prioritization axes, allowing strong candidates to emerge across different combinations of evidence rather than through a single weighted score. The resulting candidates were mapped onto a collapse-Braak target landscape and annotated by MitoCarta functional class (**Fig. 6B**).

**Figure 6.**
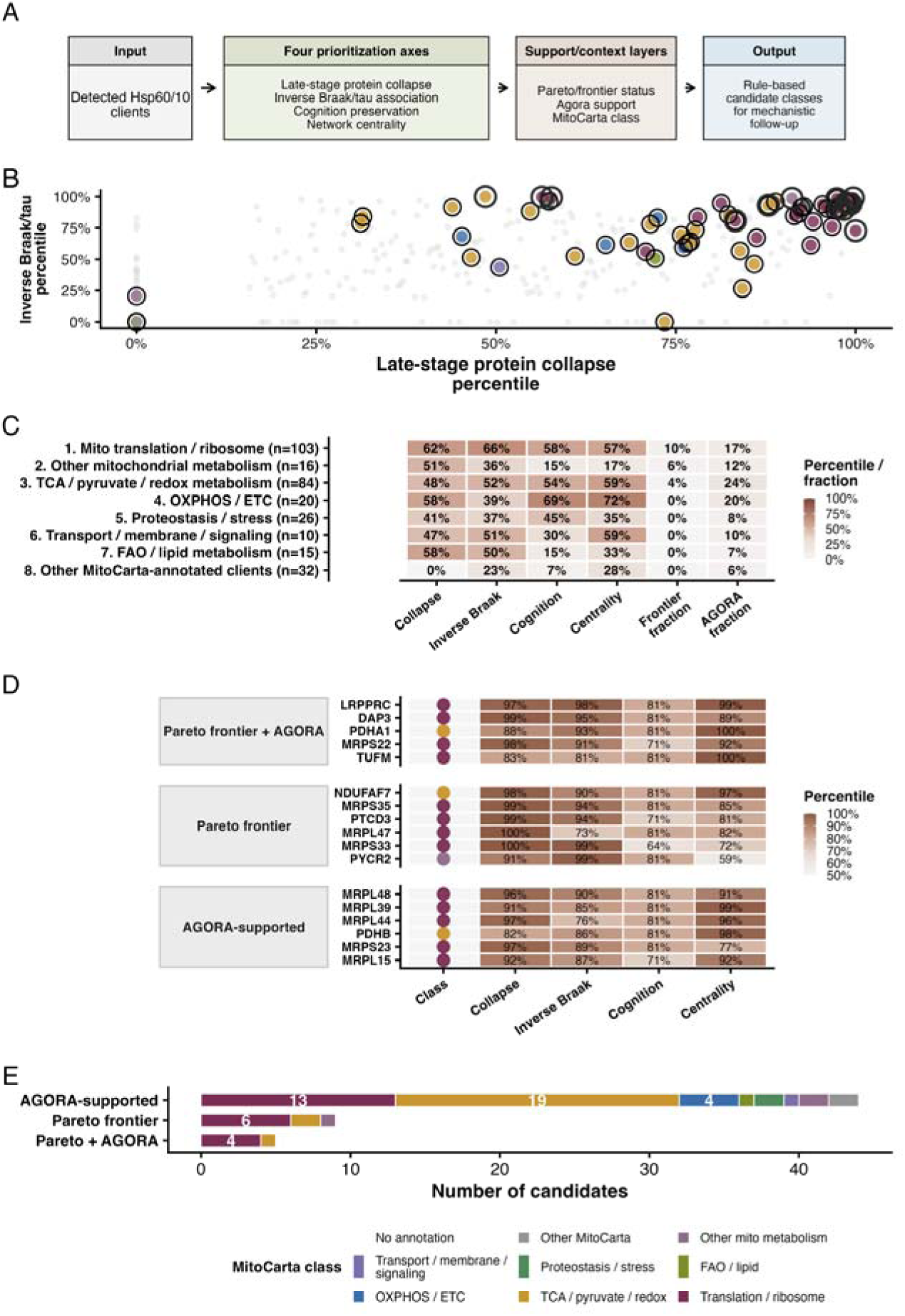
Integrated Hsp60/10 client evidence defines candidate classes for mechanistic follow-up. Detected Hsp60/10 clients (n = 306 proteins) were classified using four continuous evidence axes: late-stage protein collapse, inverse Braak/tau coupling, cognition support, and network centrality. Evidence axes were kept separate rather than collapsed into a single weighted score and were integrated with Agora nomination and MitoCarta annotation using rule-based candidate layers. (A) Candidate classification framework showing the evidence axes and non-weighted integration used to define Pareto/frontier candidates, Agora-supported candidates, and MitoCarta-annotated follow-up candidates. (B) Collapse-Braak target landscape. Each point represents a detected client, plotted by late-stage collapse percentile and inverse Braak/tau association percentile. Background clients are gray; candidate-layer genes are colored by MitoCarta class. Pareto/frontier candidates are ringed and were defined using collapse, inverse Braak/tau, cognition, and centrality evidence, not the two plotted axes alone. (C) Ranked MitoCarta functional-class evidence summary. Rows represent functional classes ordered by frontier fractions and supporting evidence. Columns show class-level summaries for collapse, inverse Braak/tau, cognition, centrality, frontier fraction, and Agora fraction. (D) Rule-based candidate-layer evidence table showing up to six representative genes per layer, with class annotation and percentile support for each evidence axis. (E) Candidate-layer composition by MitoCarta functional class.

Mitochondrial translation, broad mitochondrial metabolism, and TCA/pyruvate/redox metabolism clients showed the strongest aggregate evidence across the four prioritization axes and two support layers, Pareto/frontier status and Agora nomination (**Fig. 6C**). Several Pareto/frontier candidates were also Agora-nominated, including mitochondrial translation clients TUFM, LRPPRC, and DAP3, and the TCA/pyruvate/redox client PDHA1 (**Fig. 6D**). The framework also nominated candidates not currently supported by Agora, including PTCD3 and multiple mitochondrial ribosomal proteins (MRPs), indicating that integrated client-level evidence can identify both externally supported Agora candidates and candidates newly prioritized by this framework. Together, these results prioritize mitochondrial translation and TCA/pyruvate/redox metabolism as the dominant Hsp60/10 client classes for mechanistic follow-up (**Fig. 6E**).

## DISCUSSION

### Principal finding: Hsp60/10 client vulnerability emerges across layered protein-level evidence

Integrated ROSMAP bulk RNA-seq and TMT proteomics analyses showed that AD stage-associated mitochondrial pathway remodeling was protein-biased, but not unique to Hsp60/10 clients. We therefore extended our interrogation of Hsp60/10 clients for selective AD-relevant vulnerability across multiple levels of analysis: whether their abundance declined during late-stage disease, whether the most vulnerable clients occupied central positions within the network, whether lower abundance was more strongly associated with Braak/tau pathology or CERAD/amyloid pathology, and whether lower client abundance correlated with greater cognitive impairment. Across these layers, late-stage protein collapse identified a network-central subnetwork of clients, while pathology and cognition analyses linked lower client abundance to greater Braak/tau burden and cognitive impairment. Abundance-matched mitochondrial null testing further supported selective client vulnerability compared with non-client mitochondrial proteins across multiple AD-relevant axes. These evidence axes, together with Pareto/frontier candidate identification and Agora nomination, were used to prioritize clients. Mitochondrial translation and TCA/pyruvate/redox clients achieved the highest overall priority based on our candidate classification framework.

### Protein-biased remodeling supports proteomic analysis but does not establish client specificity

Broad pathway remodeling is a known feature of AD, as prior studies have reported that protein changes exceeded RNA changes across multiomics analyses of ROSMAP AD datasets [9]. This RNA-protein discordance may be particularly relevant for Hsp60/10 clients because chaperonin dependence is primarily post-translational: client abundance reflects not only transcript availability but also folding efficiency and protein turnover. Therefore, we did not expect Hsp60/10 clients to be uniquely altered at the pathway level. Instead, we hypothesized that client-specific vulnerability would become apparent only when client abundance was evaluated against abundance-matched non-client mitochondrial proteins across several disease-relevant axes, including late-stage collapse, pathology association, cognitive status, and Agora nomination. Importantly, the client-enrichment signal persisted after accounting for both protein abundance and mitochondrial functional module. This argues against a simple explanation in which the Hsp60/10 client signal merely reflects overrepresentation of clients within broadly vulnerable mitochondrial translation or metabolic pathways.

### Client heterogeneity nominates a network-central, collapse-prone subset

The top-collapsed quartile client candidate proteins were found to be more central to the network, indicating that this heterogeneity is not randomly distributed across the client network. Collapse-prone clients suggest the existence of a more vulnerable, collapse-prone, network-embedded subset of Hsp60/10 clients.

### Braak/tau coupling and cognition link client abundance to disease severity

The same subset of clients that exhibited high late-stage protein collapse and network-centrality also showed strong inverse Braak/tau association, indicating that this subset of Hsp60/10 clients is also aligned with AD progression-related features. Hsp60/10 clients showed weaker inverse CERAD/amyloid associations. Client abundance decline is correlated with increasing diagnostic severity of cognitive impairment, although whether this is causative remains unclear.

Supplemental analyses indicate regional differences in AD-associated protein collapse and inverse Braak coupling, with the STG showing greater alterations than the DLPFC. Stronger STG effects are consistent with temporal association cortex being more vulnerable to AD pathology than dorsolateral prefrontal regions across much of the disease course, although interpretation is constrained by the limited number and nature of regional samples [10–12].

### Prioritized client functional classes implicate mitochondrial translation and TCA/pyruvate/redox metabolism

Hsp60/10 client vulnerability in AD is not uniformly distributed across mitochondrial functional classes. Instead, the highest-priority clients cluster within mitochondrial translation and TCA/pyruvate/redox metabolism across multiple evidence axes. These clients sustain mitochondrial protein synthesis and matrix metabolic/redox homeostasis—processes that are tightly coupled to neuronal energy demand. Their decline with AD progression is therefore well-positioned to exacerbate bioenergetic failure and cognitive impairment for which our data show correlation. Client loss could represent a driver of disease progression, a downstream consequence of other pathogenic processes, or a partially independent axis of vulnerability. Our candidate nomination framework highlights clients at key metabolic control points as strong candidates for mechanistic interrogation. Reduced PDHA1 abundance, for example, would be expected to impair pyruvate utilization, limit TCA-cycle entry, and constrain respiratory substrate supply, thereby impacting oxidative phosphorylation [13,14]. Because Hsp60/10-dependent folding is ATP-dependent, such bioenergetic deficits could further reduce chaperone capacity, establishing a feed-forward cycle that amplifies client instability [5].

To determine whether this pattern extends across central metabolic nodes, we expanded STRING network analyses to clients in three groups: Pareto/frontier plus Agora-nominated, Pareto/frontier only, and Agora-nominated only. Across groups and nomination thresholds, TCA/pyruvate/redox enzymes consistently ranked among the most vulnerable. These included components of the pyruvate dehydrogenase complex (PDHA1, PDHB, DLAT, DLD) and the α-ketoglutarate dehydrogenase complex (OGDH, DLST, DLD) [15–18]. Notably, DLD is shared between both complexes and mediates redox-linked electron transfer required for generating reducing equivalents for oxidative phosphorylation, positioning it as a potential hub linking pyruvate entry, TCA-cycle flux, and mitochondrial redox balance [19,20].

A similar convergence was observed for mitochondrial translation clients. Five proteins—LRPPRC, DAP3, MRPS22, TUFM, and PDHA1—were both Pareto/frontier candidates and Agora-nominated and showed strong integration across late-stage protein decline, network centrality, inverse Braak association, and cognitive preservation. LRPPRC, DAP3, MRPS22, and TUFM support mitochondrial gene expression and respiratory chain biogenesis, reinforcing the concept that both protein synthesis capacity and central carbon metabolism represent coordinated points of vulnerability [21–24].

Together, these findings define a convergent mitochondrial axis—spanning translation and TCA/redox metabolism—that is selectively sensitive to Hsp60/10 client instability in AD. This allows us to prioritize the most likely candidates (e.g. PDHA1 or DLD) to test for direct causative contribution to effects on mitochondrial function, redox state, and tau-related phenotypes in appropriate wild-type and tauopathy models.

### Limitations and future directions: from human omics associations to mechanism

This study does not establish whether the observed decline in Hsp60/10 client abundance is a cause of mitochondrial dysfunction, a downstream consequence of neurodegeneration, or part of a reinforcing feedback loop during AD progression. Accordingly, candidate prioritization for mechanistic follow-up remains hypothesis-generating and does not, on its own, establish therapeutic relevance. Because the primary transcriptomic and proteomic analyses were performed at the bulk-tissue level, they cannot resolve cell type-specific vulnerability and may reflect changes in cell-state composition, disease-associated cellular remodeling, or both. Higher-resolution single-cell, spatial, and model-system studies will therefore be important for determining which neuronal and glial populations show the strongest Hsp60/10 client vulnerability, distinguishing cell-intrinsic proteostasis failure from shifts in tissue composition and guiding the selection of appropriate experimental models for mechanistic validation.

These studies provide strong justification for future experimental studies to establish causality by directly perturbing prioritized candidates and assessing effects on mitochondrial function and disease-relevant phenotypes. Beyond genetic approaches, pharmacological modulation of the Hsp60/10 system represents a complementary parallel strategy. Enhancing chaperone function could potentially restore the abundance of vulnerable, network-central clients and mitigate AD-related phenotypes, including cognitive impairment and tau pathology. Emerging evidence supports the feasibility of this approach. For example, mizoribine has been reported to stabilize the Hsp60/10 complex by preventing Hsp10 dissociation, potentially promoting client folding [25]. In parallel, unpublished work from our group has potentially identified a small molecule as a direct Hsp10-binding compound. Notably, this same small molecule reduces pathogenic tau burden in a mouse model of AD [26], suggesting that targeting the Hsp60/10 complex may represent a viable therapeutic strategy linking mitochondrial proteostasis to tau pathology.

## DECLARATIONS

### Ethics approval and consent to participate

This study involved secondary analysis of de-identified human participant data obtained from Synapse/AD Knowledge Portal under applicable data access and data use agreements. No new human participants were recruited, no new biospecimens were collected, and no identifiable participant-level information is reported in this manuscript. Ethics approval and informed consent for the original cohort studies were obtained by the respective data-generating institutions. No additional ethics approval or participant consent was sought for the present secondary analysis of de-identified data.

### Consent for publication

Not applicable.

### Availability of data and materials

The datasets analyzed during the current study are available through the AD Knowledge Portal, Synapse, Agora, and associated repositories, subject to applicable data access requirements and data use agreements.

ROSMAP data are available through the AD Knowledge Portal: https://doi.org/10.7303/9618239. MSBB data are available through the AD Knowledge Portal: https://doi.org/10.7303/9618238.

AMP-AD Diverse Cohorts data are available through the AD Knowledge Portal Diverse Cohort Study: https://doi.org/10.7303/9618093. Agora is available at: https://doi.org/10.57718/Agora-adknowledgeportal.

Derived analysis files, source data for figures, and code supporting the findings of this study will be made available upon publication in an archived analysis repository (https://doi.org/10.5281/zenodo.20851582). Protected ROSMAP, AMP-AD, Synapse-controlled, and other third-party source data are not redistributed and remain subject to their original data-access terms.

### Competing interests

The authors declare that they have no competing interests.

### Funding

This work was supported by NIH R25 AG086096. The funder had no role in the conceptualization, study design, data collection, data analysis, interpretation of results, decision to publish, or preparation of the manuscript.

### Authors’ contributions

AS conceptualized the study, curated datasets, performed analyses, generated figures, interpreted results, and drafted the manuscript. JKA supervised the study, contributed to study conceptualization and interpretation, and critically revised the manuscript. VK contributed to study conceptualization and critically revised the manuscript. All authors read and approved the final manuscript.

## Acknowledgements

The authors thank Dr. Greg Chin for his invaluable feedback on the manuscript.

## ROSMAP

The results published here are in whole or in part based on data obtained from The AD Knowledge Portal (https://doi.org/10.7303/9618239). Study data were provided by the Rush Alzheimer’s Disease Center, Rush University Medical Center, Chicago. Data collection was supported through funding by NIA grants P30AG10161 (ROS), R01AG15819 (ROSMAP; genomics and RNAseq), R01AG17917 (MAP), R01AG36836 (RNAseq), RF1AG57473 (single nucleus RNAseq), U01AG46152 (ROSMAP AMP-AD, targeted proteomics), U01AG46161 (TMT proteomics), U01AG61356 (whole genome sequencing, targeted proteomics, ROSMAP AMP-AD), P30AG072975, the Illinois Department of Public Health (ROSMAP), and the Translational Genomics Research Institute (genomic). Additional phenotypic data can be requested at www.radc.rush.edu.

## TMT Proteomics

Study data were provided through the Accelerating Medicine Partnership for AD (U01AG046161 and U01AG061357) based on samples provided by the Rush Alzheimer’s Disease Center, Rush University Medical Center, Chicago. Data collection was supported through funding by NIA grants P30AG10161, R01AG15819, R01AG17917, R01AG30146, R01AG36836, U01AG32984, U01AG46152, the Illinois Department of Public Health, and the Translational Genomics Research Institute.

## RNA-seq Bulk Brain

Annie J. Lee, Yiyi Ma, Lei Yu, Robert J. Dawe, Cristin McCabe, Konstantinos Arfanakis, Richard Mayeux, David A. Bennett, Hans-Ulrich Klein, and Philip L. De Jager. Multi-region brain transcriptomes uncover two subtypes of aging individuals with differences in Alzheimer’s disease risk and the impact of APOEε4. bioRxiv 2021.

## Single-nucleus RNA-seq Data

Study data were generated from postmortem brain tissue provided by the Religious Orders Study and Rush Memory and Aging Project (ROSMAP) cohort at Rush Alzheimer’s Disease Center, Rush University Medical Center, Chicago. This work was funded by NIH grants U01AG061356 (De Jager/Bennett), RF1AG057473 (De Jager/Bennett), and U01AG046152 (De Jager/Bennett) as part of the AMP-AD consortium, as well as NIH grants R01AG066831 (Menon) and U01AG072572 (De Jager/St George-Hyslop).

## MSBB

The results published here are in whole or in part based on data obtained from The AD Knowledge Portal (https://doi.org/10.7303/9618238). These data were generated from postmortem brain tissue collected through the Mount Sinai VA Medical Center Brain Bank and were provided by Dr. Eric Schadt from Mount Sinai School of Medicine.

## MSBB Proteomics

The results published here are in whole or in part based on data obtained from The AD Knowledge Portal (https://doi.org/10.7303/9618238). These data were provided by Dr. Levey from Emory University based on postmortem brain tissue collected through the Mount Sinai VA Medical Center Brain Bank provided by Dr. Eric Schadt from Mount Sinai School of Medicine.

## AMP-AD Diverse Cohorts

The results published here are in whole or in part based on data obtained from the AD Knowledge Portal Diverse Cohort Study DOI (https://doi.org/10.7303/9618093). Data generation was supported by the following NIH grants: U01AG046139, U01AG046170, U01AG061357, U01AG061356, U01AG061359, and R01AG067025. We thank the participants of the Religious Order Study, Memory and Aging Project, the Minority Aging Research Study, Rush Alzheimer’s Disease Research Center, Mount Sinai/JJ Peters VA Medical Center NIH Brain and Tissue Repository, National Institute of Mental Health Human Brain Collection Core (NIMH HBCC), Mayo Clinic Brain Bank, Sun Health Research Institute Brain and Body Donation Program, Goizueta Alzheimer’s Disease Research Center, New York Brain Bank at Columbia University, New York Genome Center and the Biggs Institute Brain Bank for their generous donations. Data and analysis contributing investigators include Nilüfer Ertekin-Taner, Minerva Carrasquillo, Mariet Allen (Mayo Clinic, Jacksonville, FL), David Bennett, Lisa Barnes (Rush University), Philip De Jager, Vilas Menon (Columbia University), Bin Zhang, Vahram Haroutanian (Icahn School of Medicine at Mount Sinai), Allan Levey, Nick Seyfried (Emory University), Rima Kaddurah-Daouk (Duke University), Steve Finkbeiner (University of California-San Francisco/Gladstone Institutes), Daifeng Wang (University of Wisconsin-Madison), Stefano Marenco (NIMH HBCC), Anna Greenwood, Abby Vander Linden, Laura Heath, William Poehlman (Sage Bionetworks).

## Agora

The results published here are in whole or in part based on data obtained from Agora, a platform initially developed by the NIA-funded AMP-AD consortium that shares evidence in support of AD target discovery. Agora is available at: https://doi.org/10.57718/Agora-adknowledgeportal.

## List of Abbreviations

AD: Alzheimer’s disease
AMP-AD: Accelerating Medicines Partnership–Alzheimer’s Disease
APOEε4: apolipoprotein E ε4 allele
AsymAD: asymptomatic Alzheimer’s disease
ATP: adenosine triphosphate
CERAD: Consortium to Establish a Registry for Alzheimer’s Disease
cogdx: Final Consensus Cognitive Diagnosis
DESeq2: R/Bioconductor package for differential expression analysis of sequencing count data
DLPFC: dorsolateral prefrontal cortex
ETC: electron transport chain
FAO: fatty-acid oxidation
FDR: false discovery rate
HSP: heat shock protein
Hsp10: heat shock protein 10
Hsp60: heat shock protein 60
Hsp60/10: Hsp60/Hsp10 mitochondrial chaperonin complex
Hsp70: heat shock protein 70
Hsp90: heat shock protein 90
MAP: Memory and Aging Project
MCI: mild cognitive impairment
MMSE: Mini-Mental State Examination
MRP: mitochondrial ribosomal protein
MRPs: mitochondrial ribosomal proteins
MSBB: Mount Sinai Brain Bank
NCI: no cognitive impairment
NIA: National Institute on Aging
NIH: National Institutes of Health
NIMH: National Institute of Mental Health
OXPHOS: oxidative phosphorylation
OXPHOS/ETC: oxidative phosphorylation/electron transport chain
PMI: postmortem interval
pQTL: protein quantitative trait locus
RIN: RNA integrity number
RNA: ribonucleic acid
RNA-seq: RNA sequencing
ROSMAP: Religious Orders Study and Memory and Aging Project
SEM: standard error of the mean
snRNA-seq: single-nucleus RNA sequencing
STG: superior temporal gyrus
STRING: Search Tool for the Retrieval of Interacting Genes/Proteins
TCA: tricarboxylic acid
TMT: tandem mass tag
TREAT-AD: Target Enablement to Accelerate Therapy Development for Alzheimer’s Disease
UPRmt: mitochondrial unfolded protein response
VA: Veterans Affairs
VST: variance-stabilizing transformation

## FIGURES

**Supplemental Figure 1.**
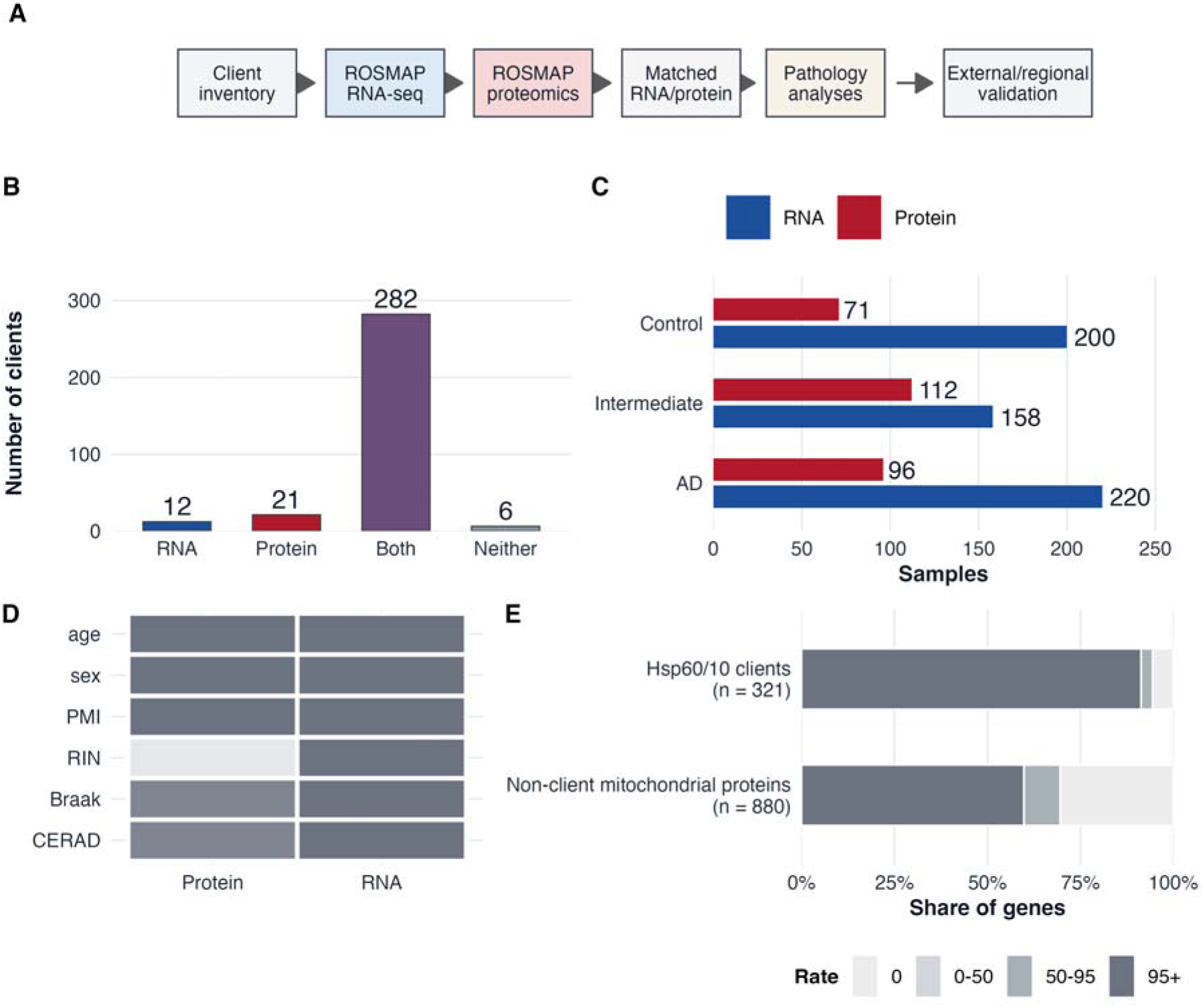
Cohort, detection, and Hsp60/10 client-set definition. Summary of the analytical workflow, Hsp60/10 client detection across ROSMAP RNA-seq and TMT proteomics, diagnostic-stage sample composition, covariate availability, and protein detection-rate categories. (A) Analysis workflow integrating a published Hsp60/10 client inventory with ROSMAP RNA-seq, TMT proteomics, matched RNA/protein subsets, pathology analyses, and external or regional validation analyses. (B) Detection of the 321-client reference inventory across RNA-seq and proteomics: RNA only (n = 12 clients), protein only (n = 21), both RNA and protein (n = 282), and neither (n = 6). (C) RNA-seq and TMT proteomics sample counts across NCI, MCI, and AD diagnostic-stage groups: RNA n = 200, 158, and 220 and protein n = 71, 112, and 96 for NCI, MCI, and AD respectively. (D) Availability of covariates and pathology variables used across RNA and protein analyses. (E) Protein detection-rate categories for Hsp60/10 clients (n = 321) and non-client mitochondrial proteins (n = 880). Detection-rate bins were 0%, >0–50%, 50–95%, and ≥95%.

**Supplemental Figure 2.**
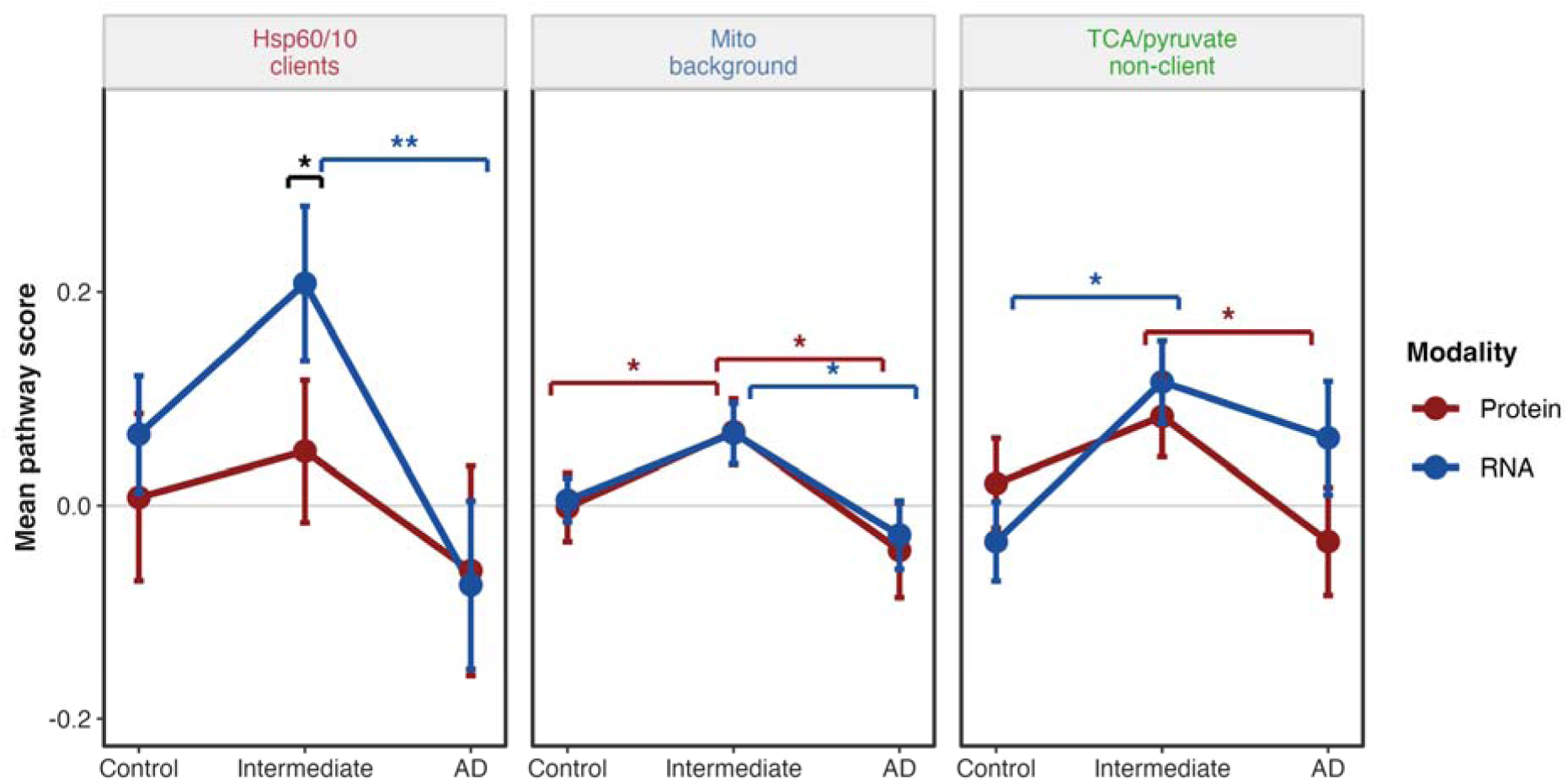
Matched-individual RNA/protein sensitivity analysis. Matched ROSMAP samples with both RNA-seq and TMT proteomics were used to test whether stage-associated remodeling patterns were preserved when RNA and protein were compared within the same individuals. Mean standardized pathway scores are shown across NCI, MCI, and AD groups for the detected Hsp60/10 client network, broad mitochondrial background, and TCA/pyruvate non-Hsp60/10 comparator pathway. Red indicates protein and blue indicates RNA. Points show group means; error bars show SEM. Colored brackets indicate within-modality stage comparisons. No significant protein-versus-RNA differences were detected within the same stage. *P < 0.05, **P < 0.01, ***P < 0.001.

**Supplemental Figure 3.**
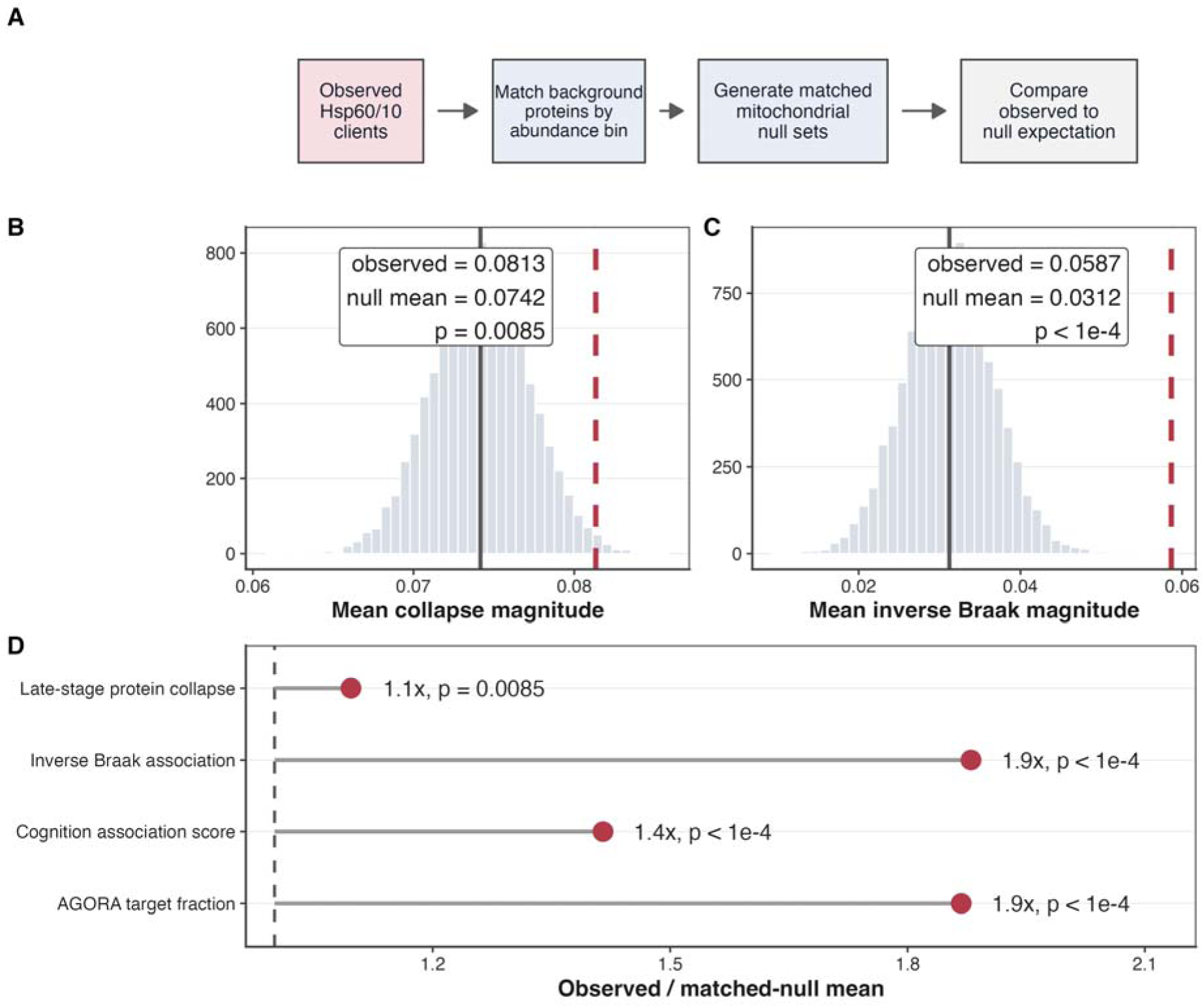
Specificity against non-client mitochondrial proteins. Observed Hsp60/10 client metrics were compared with 10,000 abundance-matched non-client mitochondrial null sets to test whether client-network signals exceeded matched mitochondrial background expectations. Values greater than 1 indicate stronger observed Hsp60/10 signal than matched-null expectation. The observed set contained n = 306 detected Hsp60/10 clients, and the matched pool contained n = 609 non-client mitochondrial proteins. (A) Schematic of the abundance-matched null strategy. (B) Null distribution of mean late-stage protein collapse magnitude: observed = 0.0813, null mean = 0.0742, empirical P = 0.0085. The dashed red line marks the observed Hsp60/10 client mean; the gray line marks the null mean. (C) Null distribution of mean inverse Braak association magnitude: observed = 0.0587, null mean = 0.0312, empirical P < 1.0 x 10^−4^, with values bound by 10,000 null iterations. (D) Observed-to-null ratios were 1.1 for late-stage protein collapse, 1.9 for inverse Braak association, 1.4 for cognition association score, and 1.9 for Agora target fraction; empirical P = 0.0085 for collapse and P < 1.0 x 10^−4^, with values bound by 10,000 null iterations, for the remaining metrics.

**Supplemental Figure 4.**
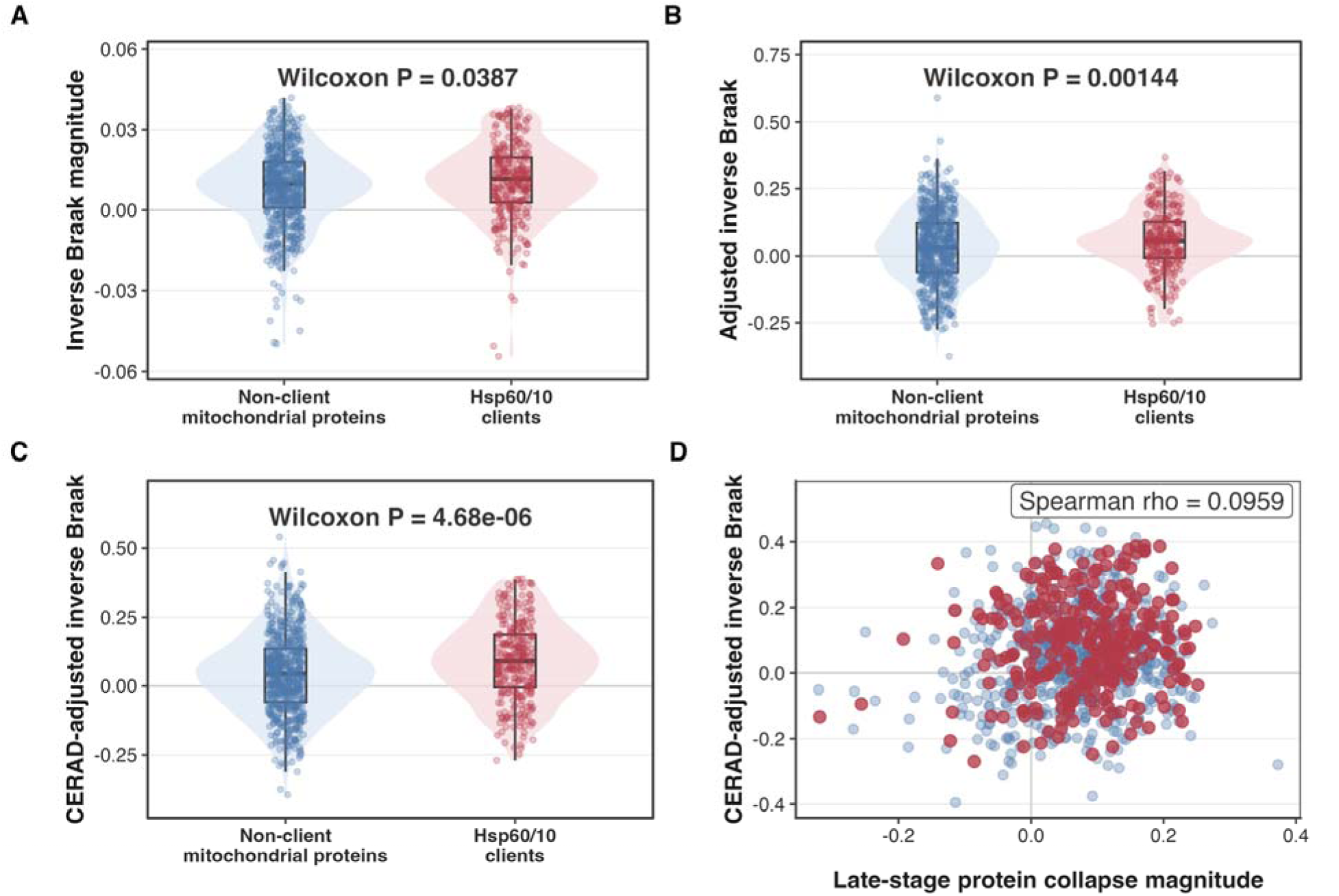
Pathology model robustness of Hsp60/10 client vulnerability. Alternative pathology models were used to test whether Hsp60/10 client vulnerability was robust to model specification. Positive inverse Braak values indicate lower protein abundance with higher Braak pathology. Hsp60/10 clients (n = 306 proteins) were compared with detected non-client mitochondrial proteins (n = 609 proteins) using two-sided Wilcoxon rank-sum tests. (A) Unadjusted inverse Braak magnitude from Spearman associations, P = 0.0387. (B) Covariate-adjusted inverse Braak beta values from models including Braak stage, age, sex, and PMI, P = 0.00144. (C) CERAD-adjusted inverse Braak beta values from models including Braak stage, CERAD score, age, sex, and PMI, P = 4.68 x 10^−6^. (D) Late-stage collapse magnitude was correlated with CERAD-adjusted inverse Braak beta across mitochondrial proteins using Spearman correlation, rho = 0.0959.

**Supplemental Figure 5.**
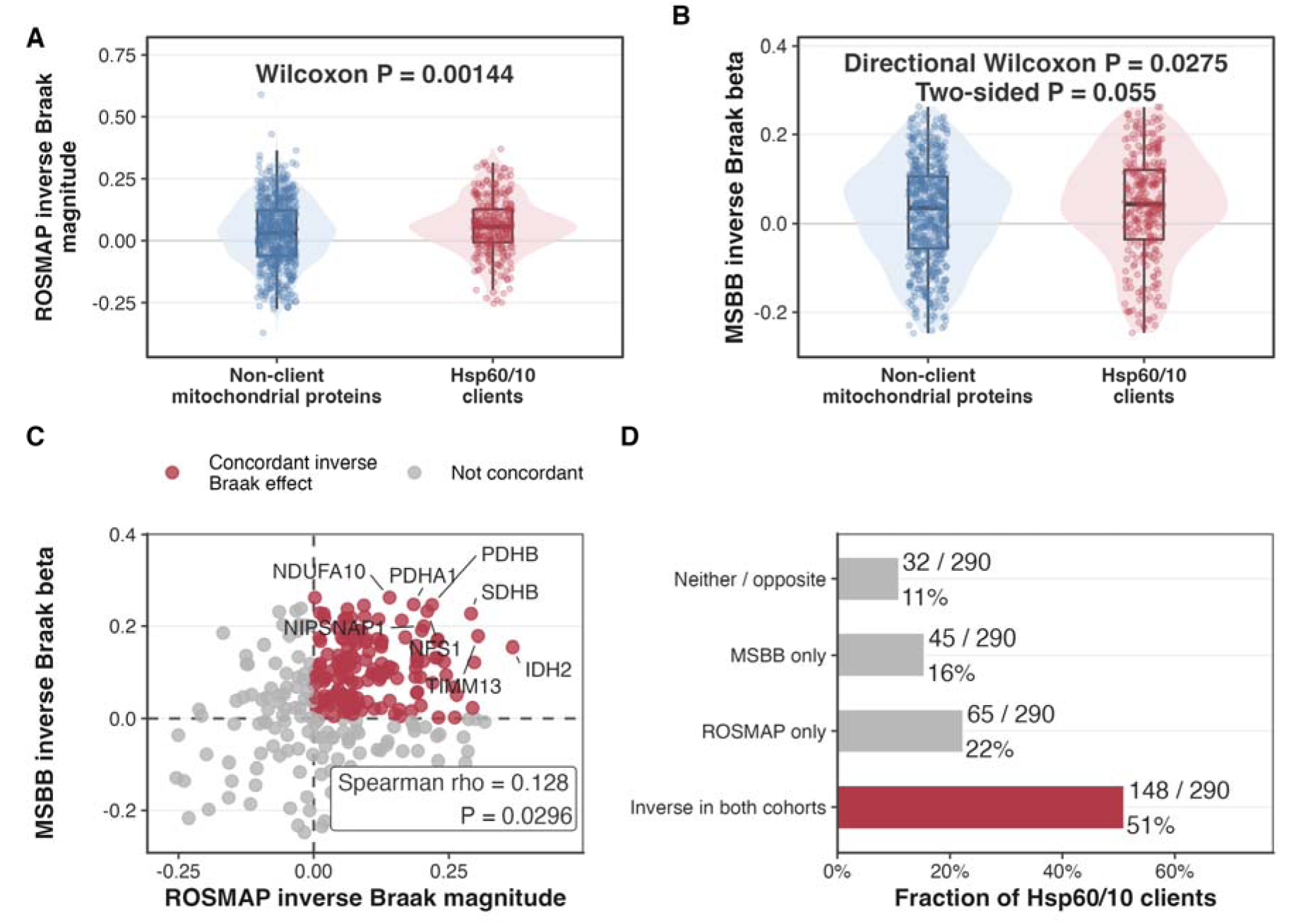
External MSBB cross-cohort validation. Independent MSBB proteomics data were used to evaluate whether Hsp60/10 client pathology coupling observed in ROSMAP was directionally reproduced across cohorts. Positive inverse Braak values indicate lower protein abundance with greater pathology burden. (A) ROSMAP inverse Braak magnitude compared between Hsp60/10 clients (n = 306 proteins) and non-client mitochondrial proteins (n = 609 proteins) using a two-sided Wilcoxon rank-sum test, P = 0.00144. (B) MSBB inverse Braak beta values compared between Hsp60/10 clients and non-client mitochondrial proteins using a two-sided Wilcoxon rank-sum test, one-sided P = 0.0275; two-sided P = 0.055. (C) Gene-level comparison of ROSMAP and MSBB inverse Braak effects for Hsp60/10 clients detected in both cohorts (n = 290 proteins) using Spearman correlation, rho = 0.128, P = 0.0296. Red points indicate concordant inverse Braak effects in both cohorts; gray points indicate non-concordant effects. Dashed lines mark zero effect. (D) Concordance categories among the n = 290 shared clients are shown as counts and percentages.

**Supplemental Figure 6.**
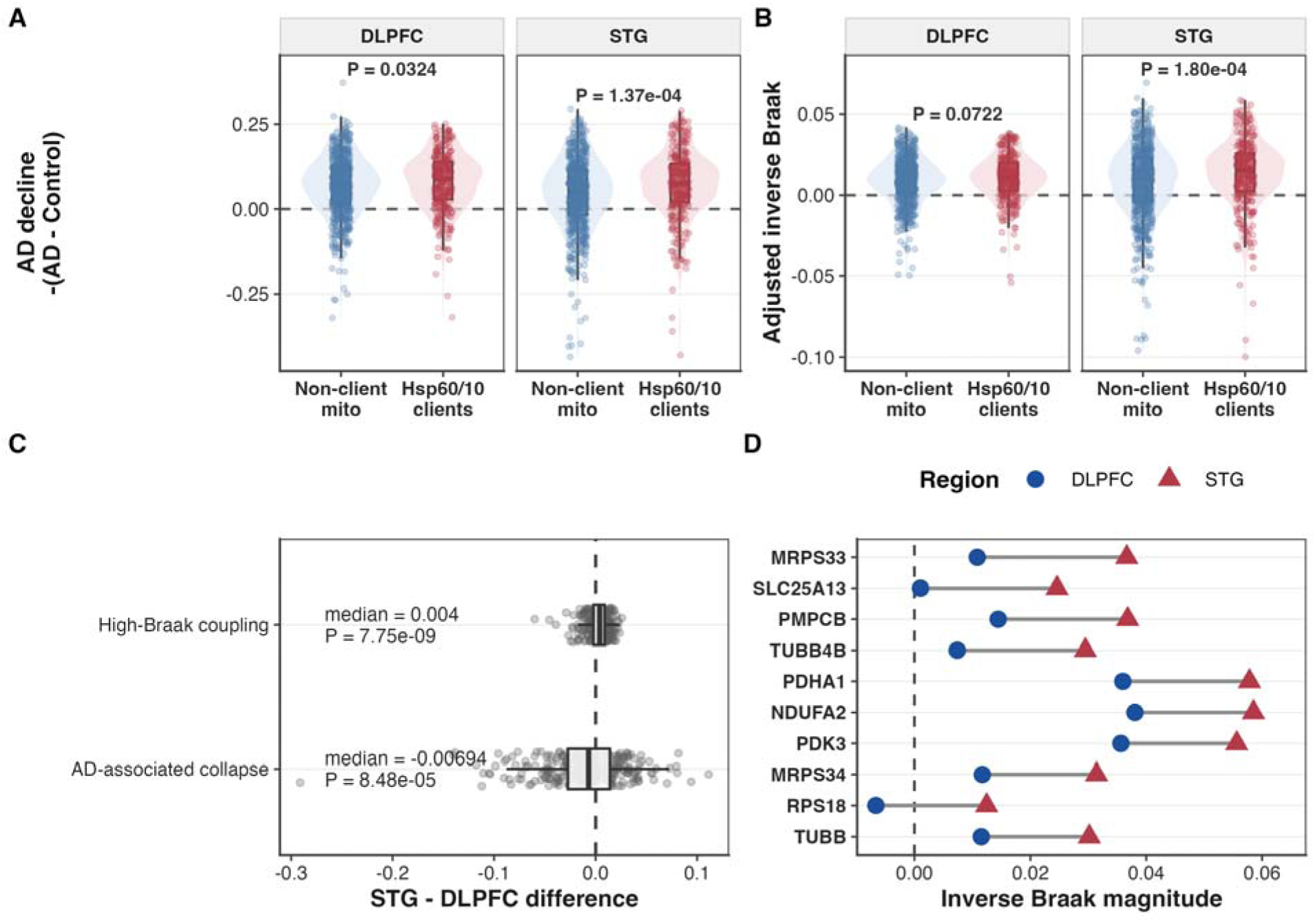
Regional proteomics validation of Hsp60/10 client vulnerability. Regional proteomics analyses in dorsolateral prefrontal cortex (DLPFC) and superior temporal gyrus (STG) tested whether Hsp60/10 clients showed regional evidence of AD– and pathology-associated protein vulnerability. Hsp60/10 clients were compared with detected non-client mitochondrial proteins. (A) AD-associated protein decline in DLPFC and STG, oriented so positive values indicate greater decline in AD versus NCI: DLPFC P = 0.0324; STG P = 1.37 x 10^−4^. (B) Adjusted inverse Braak-associated protein decline: DLPFC P = 0.0722, STG P = 1.80 x 10 ^−4^. (C) STG-minus-DLPFC differences among Hsp60/10 clients were tested using paired Wilcoxon signed-rank tests: high-Braak coupling median = 0.004, P = 7.75 x 10^−9^; AD-associated collapse median = –0.00694, P = 8.48 x 10^−5^. Positive values indicate stronger signals in STG; negative values indicate stronger signals in DLPFC. (D) Hsp60/10 clients ranked by STG-minus-DLPFC adjusted inverse Braak magnitude. Points show descriptive, region-specific values, and connecting lines show within-gene regional shifts.

## Notes

https://github.com/asloane-buck/hsp60-10_rosmap_publication_submission

https://doi.org/10.7303/9618239

https://doi.org/10.7303/9618238

https://doi.org/10.7303/9618093

https://doi.org/10.57718/Agora-adknowledgeportal

https://doi.org/10.1007/s12192-020-01080-6

### Competing Interest Statement

The authors have declared no competing interest.

